# Resolving eukaryotic river biofilm communities using long-read sequencing for biomonitoring

**DOI:** 10.64898/2026.02.20.706759

**Authors:** Meri A. J. Anderson, Amy C. Thorpe, Susheel Bhanu Busi, Jonathan Warren, Kerry Walsh, Daniel S. Read

## Abstract

Freshwater biofilms host diverse microbial eukaryotic communities that are central to ecosystem functioning and serve as key indicators of water quality. Molecular biomonitoring approaches based on environmental DNA (eDNA) sequencing are increasingly used to characterise these communities, offering scalable alternatives to traditional microscopy-based assessments. Understanding how DNA sequencing methods influence the observed community composition and diversity is essential for ensuring accurate ecological interpretation. Here, we compared short-read Illumina and long-read Pacific Biosciences sequencing of the 18S rRNA gene, alongside a trimmed long-read dataset (restricted to the Illumina-primed region), to evaluate how read length and sequencing platform affect community profiling in river biofilms from seven English rivers sampled across three timepoints.

Distinct community patterns were observed between the sequencing approaches, with PERMANOVA revealing significant differences in beta diversity (p = 0.001) and modest effect sizes (R^2^ = 3.8–8.3%). While the long and trimmed datasets produced nearly identical community structures, both diverged strongly from the short-read data, suggesting that short-read sequencing captures a systematically different subset of taxa than long-read sequencing. Long-read sequencing significantly improved taxonomic resolution of the 18S rRNA gene, particularly at the genus and species levels, enabling detection of lineages that were unresolvable in short-read data. However, comparisons of paired long- and trimmed-read ASVs indicated that trimming can increase taxonomic mismatches at finer ranks, likely due to reduced sequence length rather than sequencing platform bias.

Collectively, our results demonstrate that sequencing strategy significantly influences inferred community composition and taxonomic precision. Long-read sequencing provides a more robust representation of community diversity, whereas trimmed analyses reveal how shorter amplicons may contribute to misidentification. These findings emphasise the importance of considering read length when interpreting eDNA-based assessments using the 18S rRNA gene and support the adoption of long-read sequencing for high-resolution biomonitoring applications.

## Introduction

DNA sequencing has transformed how we study the living world, opening new opportunities for understanding biodiversity, species interactions, and ecosystem functions (Shendure *et al*., 2017; Goodwin *et al*., 2016). The application of DNA sequencing in environmental research has enabled us to characterise complex, multi-kingdom eukaryotic microbial communities that are often overlooked in bacterial-centric studies (Thompson *et al*., 2017). By revealing the breadth of eukaryotic diversity in environmental samples, sequencing can assist ecologists, conservationists, and policymakers in monitoring ecosystem health, detecting rare or cryptic taxa, and guiding the management of biodiversity loss (Taberlet *et al*., 2012; Porter & Hajibabaei, 2018).

One commonly used marker for eukaryotic microorganisms is the 18S ribosomal RNA (rRNA) gene, which offers a means to survey microeukaryotes across many lineages (protists, fungi, and microalgae) in freshwater and other environments. Its utility depends heavily on the choice of primer, fragment length, and sequencing platform. Several recent studies have evaluated 18S primer sets and markers. Zheng *et al*. (2022) compared 18S primers for detecting amoebae and revealed significant variations in detection depending on the primer design. Similarly, a 2024 in silico study assessed various short 18S rDNA markers (including those from global initiatives such as Tara Oceans), comparing taxonomic coverage and resolution among eight candidate markers, some of which are longer fragments, emphasising that marker length and region can strongly affect outcomes (Zimmermann *et al*., 2024).

Sequencing platforms differ in primer region, read lengths, error profiles, throughput, and cost, all of which influence their ability to capture the diversity of eukaryotic communities. Short-read platforms (e.g. MiSeq and NextSeq (Illumina), Ion Torrent (Thermo Fisher Scientific), Onso (PacBio), DNBSEQ (MGI), AVITI (Element Bioscience)) remain widely used because of their high accuracy, cost efficiency, and high throughput for detecting abundant taxa across many samples (Bentley *et al*., 2018; Satam *et al*., 2023). However, the relatively short fragments they generate may limit taxonomic resolution, particularly among closely related taxa, and may fail to differentiate some protist or small algal lineages (van Dijk *et al*., 2018). Long-read platforms (e.g. PacBio, or Oxford Nanopore) can generate much longer amplicons spanning multiple hypervariable regions, potentially improving assignment accuracy and the ability to resolve cryptic taxa (Rhoads & Au, 2015; Callahan *et al*., 2019; Logsdon *et al*., 2020). Although Chwalińska et al. (2025) demonstrated the potential of long-read 18S sequencing for enhanced protist detection relative to short reads in mixed communities and mock assemblages, its implications for downstream ecological interpretation in complex, natural biofilm environments remain underexplored. In fungal community studies, long-read metabarcoding has revealed distinct communities and sometimes lower per-sample diversity compared to short-reads, likely due to read quality, error, or amplification biases (Mittelstrass *et al*., 2025).

Biofilms are habitats where eukaryotic microorganisms are central to community structure and ecosystem functioning, such as primary production, organic matter breakdown and nutrient cycling. High-throughput sequencing of the 18S rRNA gene has revealed that freshwater biofilms harbour an exceptionally rich and taxonomically diverse assemblage of microbial eukaryotes, including diatoms, green algae, fungi, ciliates, and other protists. These communities are often highly heterogeneous and include many rare or cryptic taxa that are difficult to detect using traditional microscopy-based approaches. As a result, 18S rRNA metabarcoding has become an increasingly important tool for capturing the full breadth of eukaryotic diversity in biofilm systems and for improving ecological interpretation of freshwater environments. Recent studies applying 18S sequencing to biofilms across a range of substrates and ecosystems have consistently demonstrated the presence of complex and previously underappreciated eukaryotic assemblages. For example, Reisoglu et al. (2024) showed that biofilms on both marine and freshwater microplastics contained diverse microalgal, fungal, and protistan communities when analysed using 18S rRNA data. Such findings highlight the ability of 18S metabarcoding to uncover substantial hidden diversity within biofilms and to resolve community components that are central to ecosystem functioning but remain poorly characterised.

Despite these advances, relatively few studies have systematically compared short- and long-read sequencing approaches for 18S rRNA gene amplicons in freshwater biofilms. The extent to which long-read sequencing improves fine-scale taxonomic resolution, enhances detection of rare taxa, and alters community composition relative to short-read methods remains insufficiently characterised. In addition, the trade-offs among primer choice, sequencing coverage, read length, cost, throughput, and error rates are not yet fully understood within natural freshwater biofilm systems.

In this study, we addressed these gaps by comparing short-read Illumina and long-read PacBio sequencing of the 18S rRNA gene from epilithic river biofilm samples collected from seven sites across England (42 samples). We tested the hypothesis that long-read sequencing in biofilms would provide higher species-level resolution of eukaryotic microflora, reveal taxa that are unassigned in short-read datasets, and more effectively capture rare or low-abundance eukaryotic taxa. By characterising both overlapping and unique detections and comparing community composition and diversity metrics across platforms, we aim to better understand the methodological trade-offs relevant to environmental biomonitoring.

## Methods

### Sample Collection

Epilithic biofilm samples were collected from rivers across England as part of the Environment Agency’s River Surveillance Network (RSN) monitoring program, following the standard sampling method described by Kelly *et al*. (2020). The total sampling campaign encompassed 2,101 river biofilm samples collected from 861 sites between 2021 and 2023. A detailed description of the methods for sample collection and short-read sequencing is described elsewhere (Environment Agency, 2024; Thorpe et al. 2025). This study focused on a subset of 42 samples collected from seven sites, each sampled twice per year in 2021, 2022, and 2023 (Supplementary Figure 2). The 2021 and 2022 UK Centre for Ecology & Hydrology (UKCEH) land cover maps (Morton *et al*., 2021) were used to describe the dominant land cover types in the upstream catchment of each site (Supplementary Table 3). Samples were obtained by scraping biofilm from five stones or macrophytes into a tray containing 50 ml of river water. The upper surfaces were brushed with a clean toothbrush to remove biomass. A 5 ml aliquot of the biofilm suspension was then transferred to a centrifuge tube using a pipette and preserved in an equal volume of DNA preservation buffer (3.5 M ammonium sulphate, 17 mM sodium citrate, and 13 mM EDTA). Following collection, samples were concentrated by centrifugation at 3000× g for 15 ± 2 min at 5 ± 2 °C, frozen, and transported on dry ice to UKCEH, Wallingford, for further analysis. Over a three-month period leading up to and including the day of biofilm collection, up to five independent measurements of water chemistry were recorded at each site, and the mean value was calculated. Parameters measured included nitrate, phosphorus, dissolved oxygen, water temperature, and pH. Full water chemistry data are available in the Supplementary Material (Supplementary Table 3, Supplementary Figure 1 and 2).

### DNA extraction

DNA was extracted from 100 µL of biofilm suspension using the Quick-DNA Fecal/Soil Microbe Kit (Zymo Research, California, U.S.) with modifications to optimise the DNA yield (Newbold *et al*., 2025). The extraction protocol was adapted as follows: 500 µL of DNA/RNA Shield (Zymo Research) was added to each sample as a lysis buffer. The samples were mechanically disrupted using a TissueLyser II (Qiagen, Germany) at 20 Hz for 20 min, and 20 µL of recombinant Proteinase K (Roche, Switzerland) was added to the lysate and incubated at 65°C for 20 min. The purified DNA was eluted in 100 µL elution buffer. A negative extraction control was used to monitor potential contamination. The DNA concentration was quantified using the Qubit dsDNA High Sensitivity kit (Life Technologies Limited) according to the manufacturer’s protocol. The extracted DNA was stored at 4 °C until PCR amplification. A detailed step-by-step extraction, PCR, and library preparation protocol is available at https://doi.org/10.17504/protocols.io.j8nlk8em6l5r/v1.

### Sequencing

Two sequencing methods were used to amplify the 18S rRNA gene (Figure 1). Short-read sequencing (330 bp) was performed on the Illumina NextSeq platform, and long-read sequencing (1800 bp) was performed on the Pacific Biosciences Sequel II platform.

**Figure 1.**
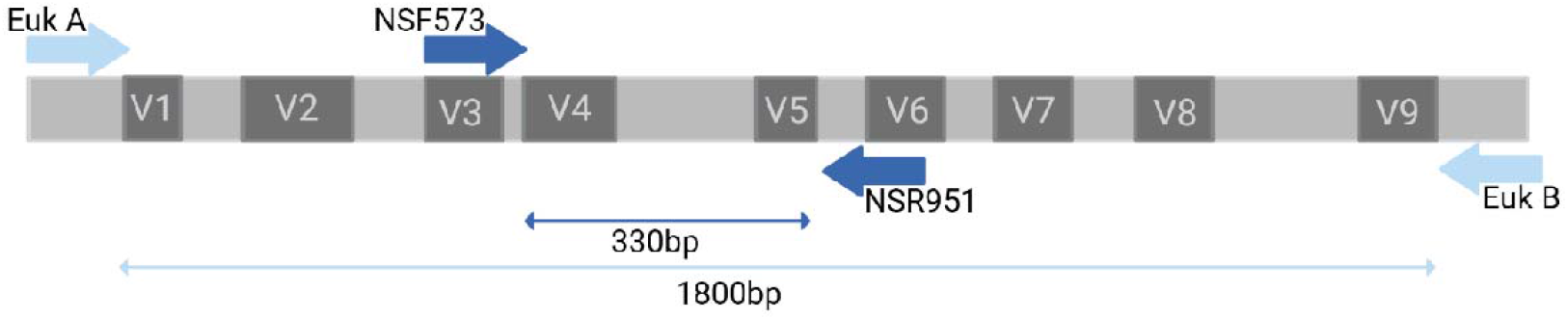
Primer positions on the 18S rRNA gene, showing the overlap of the Illumina 18S V4-V5 primers within the 18S gene sequenced by the PacBio 18s V1-V9 primers.

### Illumina

The V4-V5 region of the 18S rRNA gene was amplified using specific primers (Supplementary Table 1) modified to include Illumina adaptor sequences. In a UV-sterilised laminar flow hood, a master mix was prepared containing 0.5 µL of 2000 units mL^−1^ Q5 high-fidelity DNA polymerase, 10 µL of 5x reaction buffer, 10 µL of 5x high GC enhancer (New England Biolabs, UK), 1 µL of a 10 mM dNTP mix (Bioline, UK), 0.1 µL of each 100 µM forward and reverse primer pair (Supplementary Table 1), and 26.3 µL of molecular grade water. The master mix (48 µL) was dispensed into each well of a 96-well plate, and 2 µL of template DNA was added per sample. Negative PCR controls were also included. The thermocycling conditions are presented in Supplementary Table 2. Successful amplification was verified by 1.5% agarose gel electrophoresis using GelRed nucleic acid stain. PCR products were purified using a MultiScreen PCR filter plate, resulting in 35 µL of eluted product.

The second PCR step employed a dual-indexing approach to enable sample demultiplexing. Indexing primers were prepared using an Opentrons liquid-handling robot, each consisting of a forward (i5) or reverse (i7) Illumina adaptor sequence, i5 or i7 Nextera index, and an Illumina pre-adaptor sequence. The second PCR mix contained 0.25 µL of Q5 DNA polymerase, 5 µL of reaction buffer, 5 µL of high GC enhancer, 0.5 µL of dNTPs, 5 µL of the indexing primers (pre-prepared in the plate), 7.25 µL of molecular grade water, and 2 µL of purified PCR product from the first PCR step. The cycling protocol is presented in Supplementary Table 2. Amplification was confirmed by agarose gel electrophoresis.

The second-step PCR product was normalised using the NGS Normalization kit (Norgen Biotek, Canada) to achieve a concentration of approximately 5 ng µL^−1^. The samples were pooled by plate and quantified using a Qubit High-Sensitivity Assay Kit. The amplicon library was prepared by diluting and pooling the samples, followed by concentration and purification using the MinElute gel extraction kit (Qiagen, Germany). The final libraries were quantified, diluted to 1000 pM, and sent to The James Hutton Institute for sequencing on a NextSeq 2000 with a P1 2×300 bp flow cell and 30% PhiX control.

### Pacific Biosciences

Extracted DNA was sent to Novogene UK for PCR amplification of the V1-V9 region of the 18S rRNA gene (primer sequences and PCR amplification conditions are provided in Supplementary Table 1 and 2, respectively). Barcoded V1–V9 PCR product was verified by agarose gel electrophoresis prior to sequencing on a PacBio Sequel II sequencing platform (Novogene, Cambridge, UK).

### Data Analysis

Amplicon sequence reads were processed using the DADA2 pipeline (Callahan *et al*., 2016) implemented in R [version 4.4.2]. Short-read and long-read sequences underwent distinct processing workflows, with full analysis scripts for short-read sequences available at https://github.com/amycthorpe/amplicon_seq_processing_biofilms. For short-read analyses, raw sequences were demultiplexed, and adaptor sequences were trimmed using the Illumina FASTQ generation pipeline. Primers were removed using the ‘trimLeft’ parameter, and the quality profiles of the forward and reverse reads were examined. Reads were truncated when quality scores fell below Q30 and filtered using stringent criteria, including the removal of reads with ambiguous bases and a maximum expected error threshold of 2. The DADA2 algorithm learned error rates from a 100 million base subset, with visualisations confirming the alignment of the estimated rates with the observed data. Reads were then dereplicated into unique sequences based on the error rate model, and the core sample inference algorithm was used to identify true sequence variants. Paired forward and reverse reads were aligned and merged, requiring a minimum of a 12-base overlap. Chimeric sequences were identified and removed, resulting in an amplicon sequence variant (ASV) abundance table. Long-read analyses followed a modified protocol (https://benjjneb.github.io/LRASManuscript/LRASms_fecal.html), with primers removed and reads trimmed to a minimum length of 1,600 bp, maximum length of 1,900 bp, expected error threshold of 2, and a quality threshold of 3. Subsequent demultiplexing generated a sequence table with sample-specific counts. Taxonomy was assigned to each ASV using the naive Bayesian classifier (Wang *et al*., 2007) with a minimum bootstrap confidence of 60 against the PR^2^ version 5.0.0 SSU reference database (Guillou *et al*., 2012) for Illumina and PacBio 18S rRNA gene sequences.

To assess potential primer bias, long-read sequences were trimmed using the specific primer sequences employed in the short-read protocol to isolate the corresponding V4-V5 amplicon region. These trimmed long-read sequences were processed using the same DADA2 pipeline as the original short-read data. All subsequent analyses were conducted on these three datasets (short, long and trimmed-long) in R, using the same workflow to ensure consistency and comparability.

The sequences were rarefied to a uniform sequencing depth by examining rarefaction curves and identifying the sequencing depth at which richness plateaued (Supplementary Figure 3). Both long- and short-read data were rarefied to 4,800 reads per sample to conserve the majority of the samples. All negative extraction and PCR controls, and a small number of samples (nine) that did not meet the rarefaction depth were removed.

Downstream analyses were performed using R [version 4.4.2]. Counts, taxonomy, and metadata files were loaded into Microeco (Liu *et al*., 2021) for processing and visualisation. To assess the differences in the number of ASVs between sequencing technologies, we used the Wilcoxon signed-rank test, followed by a linear mixed-effects model to evaluate the direction of the effect. Taxonomic assignment proportions across ranks were compared using the Mann–Whitney U-test. Differences in the relative abundance of taxa between sequencing platforms were tested using the Wilcoxon rank-sum test with Bonferroni correction for multiple comparisons.

To examine similarities in community composition across sample types and sequencing platforms, principal coordinate analysis (PCoA) based on Bray–Curtis dissimilarity at the genus level was performed, along with a Procrustes analysis to assess concordance between ordinations. Prior to analysis, ASV identifiers were replaced with their corresponding taxonomic names, from kingdom to genus levels, to ensure a meaningful comparison of community structure between sequencing methods. All statistical analyses were performed using R, with significance thresholds set at p < 0.05.

Co-occurrence networks were constructed at the ASV level. To reduce noise, ASVs with fewer than two reads on average across the samples were removed prior to analysis. Networks were inferred using the SPIEC-EASI framework (Kurtz *et al*., 2015) implemented with a graphical lasso (spiec.easi function, method = “glasso”). Parameters were set with lambda.min.ratio = 1e-2, nlambda = 20, and stability selection was performed with 50 repetitions (pulsar.params = list(rep.num = 50)). The resulting adjacency matrices were used to compute network topological properties, including the number of nodes, edges, density, clustering coefficient (transitivity), modularity, assortativity, diameter, mean path length, and maximum cliques.

To compare short- and long-read ASVs in the sequence space and identify overlapping short- and long-read ASVs, we used NCBI BLAST (Camacho *et al*., 2009), with the following parameters: ‘–id 80 –query-cover 90 –subject-cover 90 –more-sensitive –outfmt 100’. The output was filtered to retain sequences with at least 90% identity and a minimum length of 300 bp for subsequent analyses. This length threshold was chosen because it represents 10% below the median short-read length. Short-read ASVs that did not map to long-read ASVs were considered unique sequences.

The relevant code for the analysis, figures, and raw data can be found at 10.5281/zenodo.18671451 and http://www.ncbi.nlm.nih.gov/bioproject/1424964 project: PRJNA1424964.

## Results

### River Surveillance Network biofilms

At each sampling point (Supplementary Figure 2a), epilithic biofilms were collected to assess bacterial biodiversity, complemented by the concurrent collection of relevant water chemistry and nutrient data (Supplementary Figure 2b–e) to understand how biotic and abiotic factors shape the bacterial community composition within the RSN. Water chemistry measurements, including pH, conductivity, and nutrient concentrations (e.g. nitrate and phosphate), varied across sites, with each parameter displaying a broad distribution (Supplementary Table 3). However, the potential physicochemical drivers of sequencing technology differences were not further explored because of the limited number of sites (n = 7), which constrained our ability to draw statistical inferences.

### Taxonomic compositions

To compare the two sequencing approaches (short Illumina vs. long PacBio reads) and the effect of trimming long reads, we assessed community composition and the relative abundance of assigned taxa. The relative read abundance differed markedly between the sequencing approaches at the division level (Figure 2). Long-read and trimmed-read datasets showed broadly similar community profiles, dominated by Opisthokonta, Stramenopiles, Rhodophyta, and Alveolata, with additional representation from Chlorophyta, Streptophyta, and Rhizaria (Supplementary Table 5). In contrast, the short-read dataset exhibited a comparatively narrower profile, with a reduced representation of several eukaryotic algal and protist lineages, including Rhodophyta (long; 12.2%, trimmed; 12%, short; 0.002%). Taxa belonging to minor divisions, such as Tubulinea, Evosea, and Actinobacteria, were detected at low relative abundance (<1%) across the datasets, although some were absent entirely from the shorter sequence profiles. The trimmed-read dataset retained most of the taxonomic diversity observed in the full-length long reads, indicating that trimming long reads with short-read primer sequences did not substantially alter higher-level taxonomic resolution.

**Figure 2.**
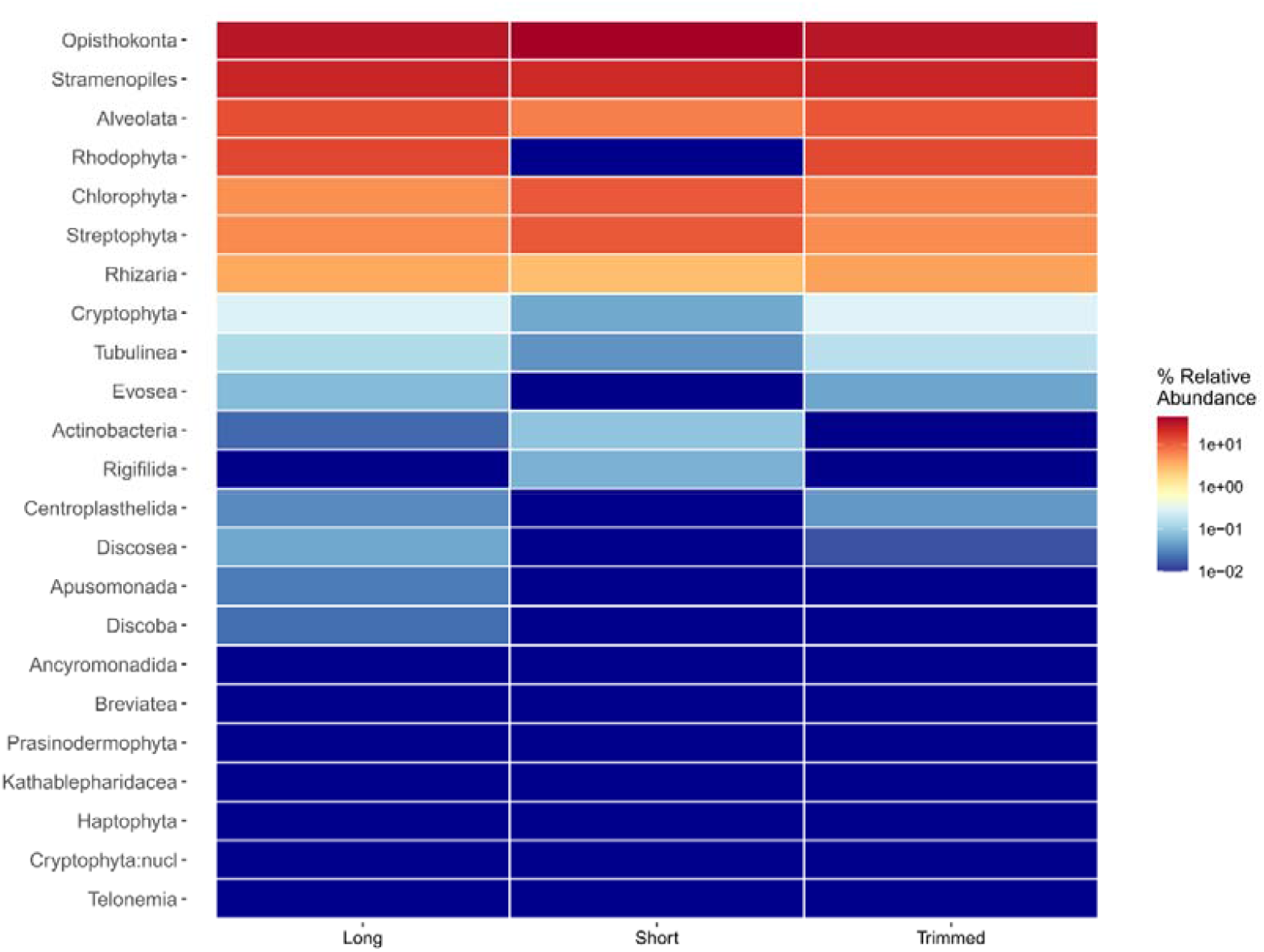
Heatmap of relative read abundance of eukaryotic Divisions detected across sequencing approaches (Long, Short, and Trimmed) based on 18S rRNA data. Colours represent log-scaled % relative abundance for each Division.

At the species level (Figure 3), distinct compositional patterns were observed between the sequencing types. Long-read sequencing recovered a broader range of taxa (114 more species in the long-read dataset, Figure 7B), including filamentous algae (*Cladophora glomerata* and *Ulothrix zonata*), diatoms (*Navicula tripunctata* and *Melosira varians*), fungi (*Hydrurus foetidus*), and metazoans (*Nais elinguis* and *Simulium santipaulii*). Short-read data captured many of the dominant species but lacked several low-abundance or rare taxa identified in the long-read and trimmed datasets, such as *Balbiania investiens, Mattesia geminata, and Polycelis tenuis* (Supplementary Table 6). The trimmed dataset showed a taxonomic profile closely aligned with the long reads, although with slightly reduced relative abundances in some groups, aligning with the short-read data. Across all sequencing types, species-level relative abundances spanned several orders of magnitude, with a few dominant taxa (S*irodotia delicatula ∼5.1%, Navicula tripunctata ∼2.3%, Brachypodium distachyon ∼2.1%*) accounting for the majority of reads and a long tail of low-abundance taxa contributing to the overall diversity.

**Figure 3.**
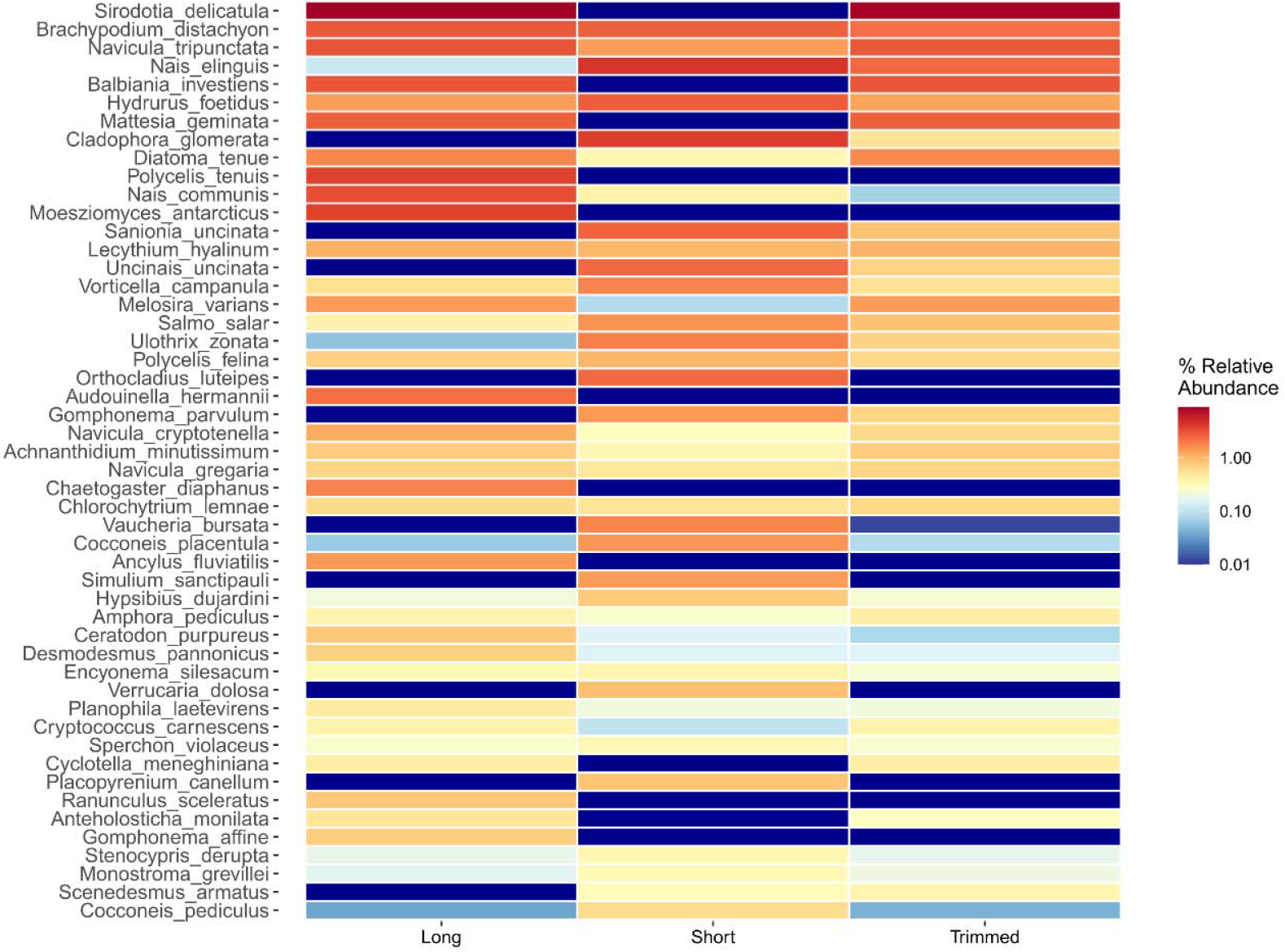
Heatmap of species-level relative abundance across sequencing approaches (Long, Short, and Trimmed) based on 18S rRNA data. Each row represents an individual species from the top 50 most abundant species across the whole dataset, with the colour intensity corresponding to the relative abundance.

Non-metric multidimensional scaling (NMDS) based on Bray–Curtis dissimilarities revealed clear separation between communities generated by short-read sequencing compared to the long-read and trimmed (Figure 4). A PERMANOVA (adonis2) indicated that sequencing type significantly influenced community composition (F = 4.07, R^2^ = 0.078, p = 0.001), with the sequencing method explaining approximately 7.8% of the total variation among samples. Pairwise PERMANOVA comparisons confirmed significant compositional differences between all methods after correction for multiple testing (Long vs. Short: F = 5.78, R^2^ = 0.083, adj. p = 0.0015; Long vs. Trimmed: F = 2.53, R^2^ = 0.038, adj. p = 0.003; Short vs. Trimmed: F = 3.85, R^2^ = 0.057, adj. p = 0.0015). Although the effect sizes were modest (R^2^ = 3.8–8.3%), each sequencing strategy captured statistically distinct communities.

**Figure 4.**
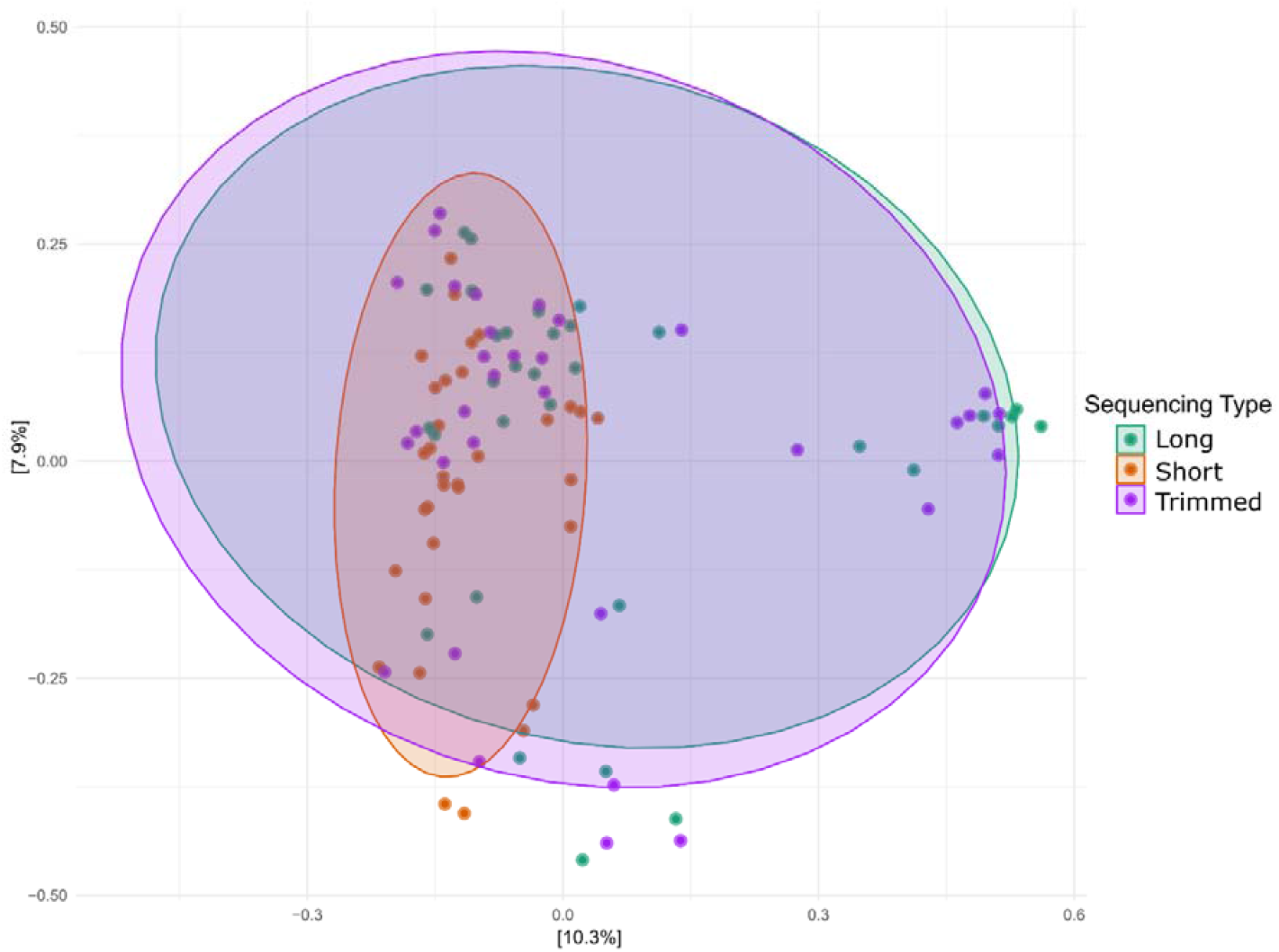
Non-metric multidimensional scaling (NMDS) ordination based on Bray–Curtis dissimilarities showing community composition across sequencing approaches (Long, Short, and Trimmed). Ellipses represent the 95% confidence intervals around the group centroids.

**Figure 5.**
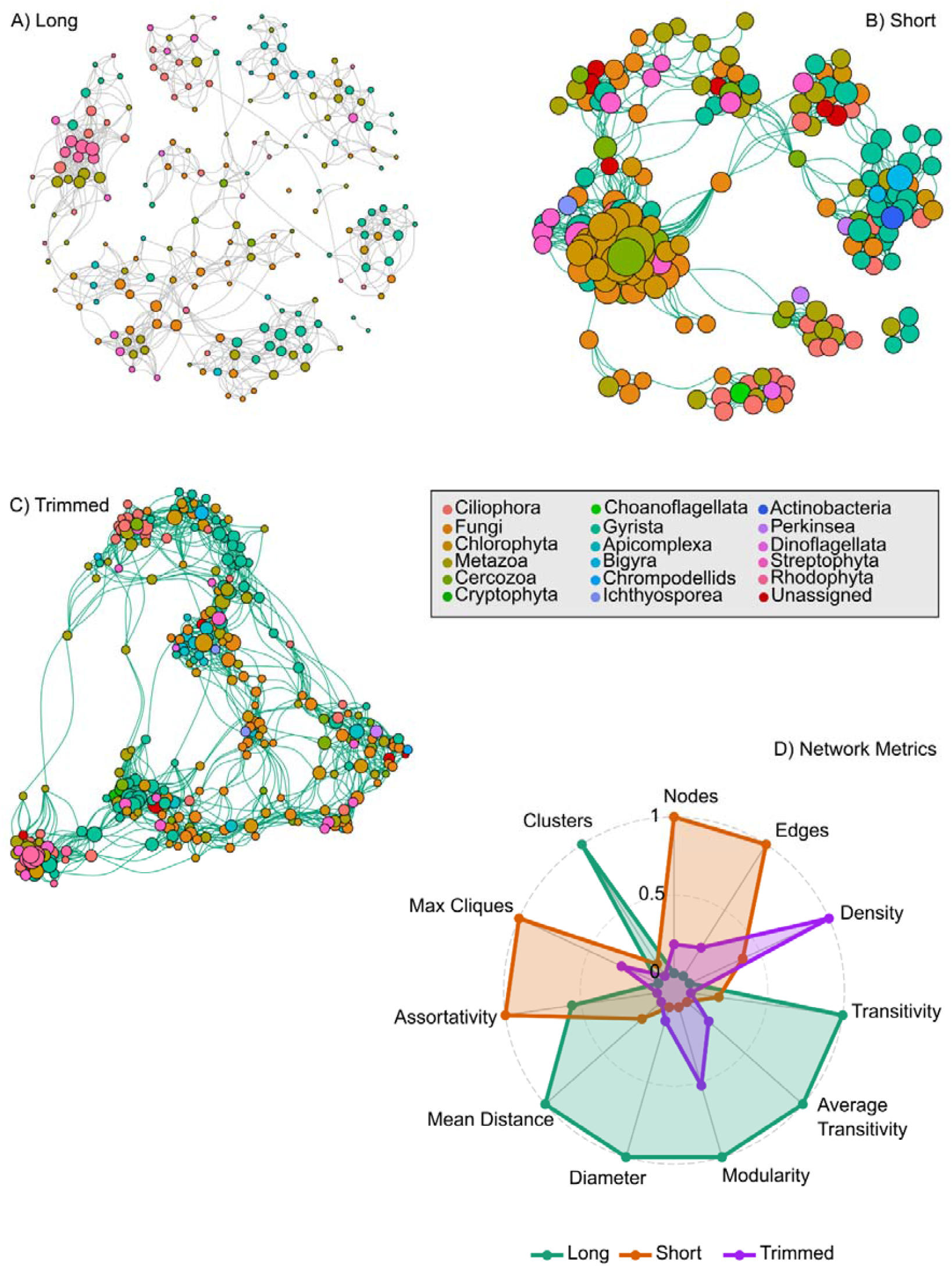
Co-occurrence network structure across long-read, short-read, and trimmed datasets. (A) Network generated from the long-read dataset. (B) Network generated from the short-read dataset. (C) Network generated from the trimmed dataset. Nodes represent ASVs and are coloured by Subdivision; node size is proportional to degree. Edge thickness represents co-occurrence strength. (D) Radar plot comparing major network metrics across the three datasets. A full summary of metrics is provided in Supplementary Table 7.

Tests for homogeneity of multivariate dispersion (betadisper) showed no evidence of unequal within-group variance between the sequencing types (F = 1.09, p = 0.325). Pairwise PERMDISP tests were all non-significant (adj. p > 0.4), indicating that the observed community differences reflected true compositional shifts rather than differences in within-group heterogeneity. Procrustes superimposition and protest tests were used to quantify concordance in community structure between sequencing approaches. Concordance between the long and trimmed datasets was very high (sum of squares = 0.0043, r = 0.998, p < 0.00001), indicating that trimming long reads with short-read primers preserved overall beta-diversity patterns. In contrast, long vs. short (sum of squares = 0.4289, r = 0.756, p = 0.60) and short vs. trimmed (sum of squares = 0.4238, r = 0.759, p = 0.55) comparisons showed only moderate, non-significant correlations, indicating weaker structural agreement. Collectively, these results demonstrate that the sequencing method significantly affects the inferred beta diversity. While all three approaches resolved distinct community profiles, the trimmed and full long-read datasets were nearly identical, suggesting that trimming long reads does not significantly distort the community structure. In contrast, short-read sequencing produced markedly different community patterns, implying potential methodological biases in primer choice and taxon detection.

### Effects of sequencing methodology on ecological networks

Network analysis was used to examine how sequencing methodology influences inferred patterns of taxon co-occurrence and ecological organisation within river biofilm communities. Co-occurrence network analysis revealed pronounced differences in network structure among the long-read, short-read, and trimmed datasets (Figure 6). Network size varied substantially across methods: the short-read dataset produced the largest and densest network (947 nodes, 8,557 edges), followed by the trimmed dataset (473 nodes, 2,687 edges), while the long-read dataset generated a comparatively smaller and more modular network (366 nodes, 1,071 edges; Supplementary Table 7). Despite its smaller size, the long-read network exhibited the highest clustering and connectivity patterns, with the greatest global transitivity (0.53), average transitivity (0.54), and modularity (0.87). In contrast, the short-read network showed lower transitivity (0.44) but a notably higher number of maximal cliques and strong assortativity (0.58), indicating strong local structuring among highly connected nodes. The trimmed dataset occupied an intermediate position between the two, showing the highest overall network density (0.024) but fewer clusters (n = 3) and reduced assortativity (0.28). Visual inspection highlighted striking differences in node prominence across sequencing methods (Figure 6A–C). Several taxa exhibited a high degree of centrality in the short-read and trimmed networks (Chlorophyta and Gyrista); however, these taxa were not consistently central in the long-read network. Instead, the long-read network distributed centrality more evenly across multiple taxonomic groups, generating a more modular and less hub-dominated architecture.

**Figure 6.**
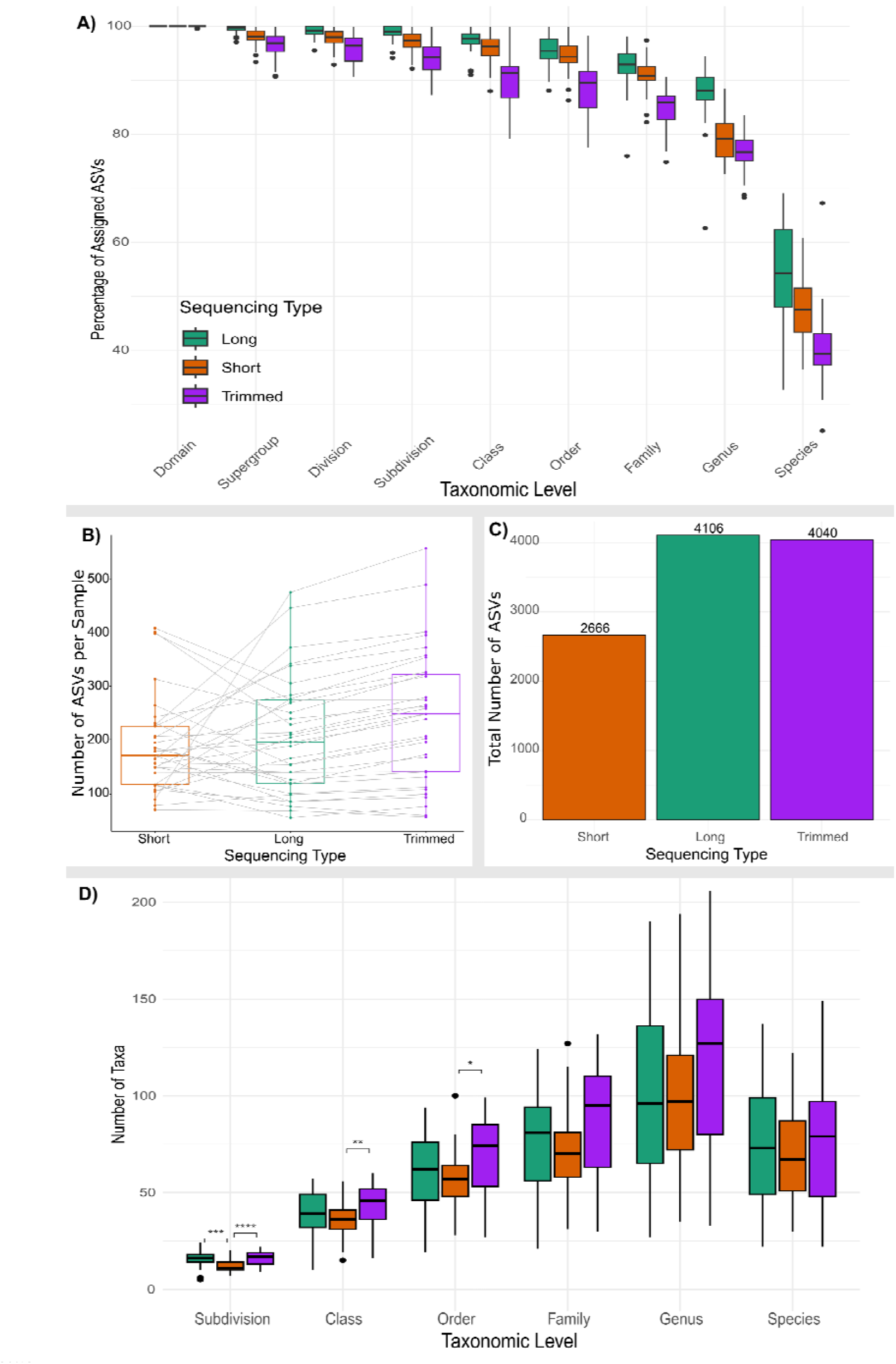
Comparison of taxonomic assignment success and richness across sequencing approaches. (A) Percentage of ASVs assigned at each taxonomic level (Domain to Species) for Long, Short, and Trimmed datasets. (B) Number of ASVs per sample by sequencing type, with grey lines connecting paired samples. (C) Total number of ASVs recovered in each dataset (Short = 2,666; Long = 4,106; Trimmed = 4,040). (D) Number of taxa identified per taxonomic level, showing significant differences among sequencing approaches (*p < 0.05; **p < 0.01; ****p < 0.0001).

**Figure 7.**
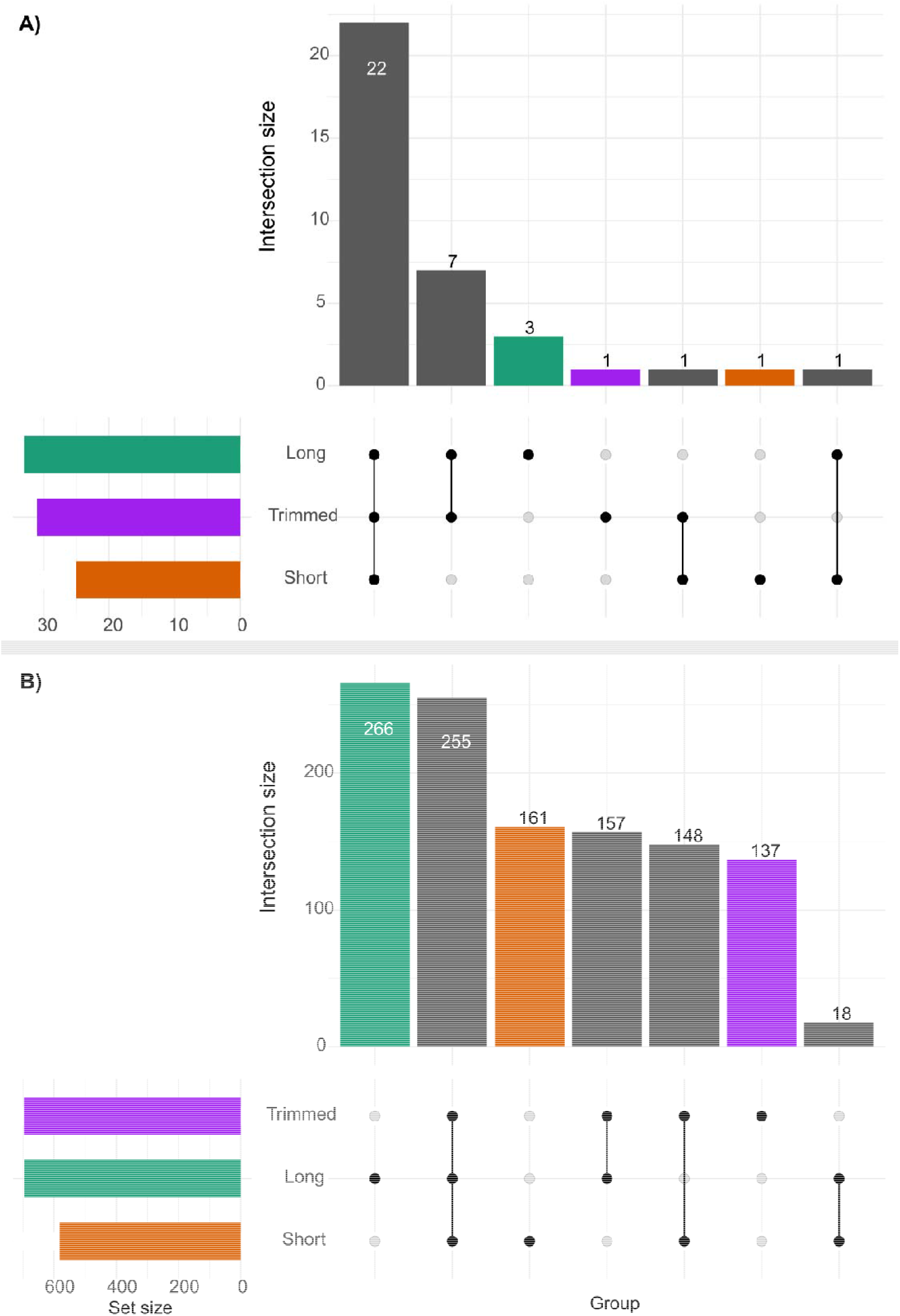
Comparison of shared taxa across the sequencing datasets. (A) UpSet plot showing the intersection of taxa detected at the subdivision level across the long-read, trimmed, and short-read datasets. (B) UpSet plot showing the intersection of taxa detected at the species level in each dataset.

Together, these results show that sequencing methodology strongly influences inferred ecological network structure, including node centrality, clustering patterns, and the extent of modular organisation. These differences emphasise the importance of sequencing approach when using network analysis as a downstream ecological inference tool.

### Improved taxonomic resolution with long reads

The proportion of ASVs successfully assigned to taxonomic levels decreased progressively from domain to species (Figure 6A). All three sequencing approaches achieved nearly complete domain-level classification (>99%), but differences became more pronounced at finer resolutions. Long-read datasets consistently achieved significantly higher assignment percentages across most taxonomic ranks than short-read and trimmed datasets (Kruskal Wallis test - Supplementary Table 4). The trimmed dataset had significantly lower taxonomic assignment than the other two methods at every taxonomic level (Supplementary Table 4). At the species level, median assignment success fell below 55% for all methods, though long reads retained slightly greater resolution.

ASV richness differed among sequencing types (Figure 6B–C). Short-read data produced the lowest ASV richness per sample (186.6 ± 88.6 ASVs), while long-read sequencing increased richness (205.3 ± 108.6 ASVs), and the trimmed dataset recovered the highest richness overall (239.7 ± 124.8 ASVs; *n* = 33 per group). A linear regression between the long-read and short-read ASV counts showed only a weak, non-significant relationship (R^2^ = 0.10, p = 0.067), and the Wilcoxon signed-rank test indicated no significant difference in their distributions (p = 0.70). In contrast, long-read counts were strongly correlated with the trimmed long-read dataset (R^2^ = 0.97, p < 0.001), although the Wilcoxon test revealed a significant difference in their overall distributions (p < 0.001), suggesting a systematic offset in absolute values despite close correspondence in relative abundance. Finally, the relationship between trimmed long-read and short-read counts was again weak and non-significant (R^2^ = 0.10, p = 0.067), with a significant difference in distributions (p = 0.046). Overall, the trimmed long-read data closely matched the untrimmed long-read counts but differed substantially from the short-read dataset, indicating that the long-read methods capture similar community patterns distinct from those observed with short-read sequencing.

In total, 2,666 ASVs were identified in the short-read dataset, compared with 4,106 in the long-read and 4,040 in the trimmed datasets, indicating that long-read sequencing recovered around 40% more unique ASVs. The trimmed dataset closely matched the long-read results, suggesting minimal information loss during trimming.

Taxonomic richness, expressed as the number of distinct taxa recovered at each rank, increased with decreasing taxonomic level (Figure 6D). Significant differences among sequencing methods were observed at several ranks, with long and trimmed reads consistently detecting more taxa than short reads (subdivision short-read – long-read: p = 0.0001 and short-read - trimmed: p = 0.00003; class short-read - trimmed: p = 0.003; order short-read - trimmed: p = 0.02). Variability among samples also increased toward the genus and species levels, reflecting the broader diversity captured by different samples.

Collectively, these results demonstrate that long-read sequencing provides deeper and more complete taxonomic resolution and richness compared with short-read sequencing, with trimming having a negligible impact on recovered diversity, but did have a significant negative impact on assigned taxonomy.

### Taxonomic overlap between each sequencing type

Comparison of taxonomic overlap between sequencing approaches showed strong concordance at the subdivision level, but substantial dissimilarity at the species level, with each dataset containing 137-266 unique species (Figure 7A–B). At the subdivision level (Figure 7A), 22 taxa were shared among all three datasets, and 7 were shared exclusively between the long- and trimmed-read datasets. At the species level (Figure 7B), 255 species were shared in the three datasets. However, 266 species were unique to the long-read dataset, 161 were unique to the short-read dataset, and 137 were unique to the trimmed dataset. The trimmed and long-read datasets shared 157 species, while the trimmed and short-read datasets shared 148 species, and only 18 species were shared exclusively between the long- and short-read datasets. These patterns indicate that the trimmed dataset largely bridges the taxonomic coverage between long- and short-read sequencing, even though each dataset still retains a substantial number of unique species.

Comparison of paired ASVs between sequencing approaches revealed differing patterns of taxonomic concordance depending on whether long-read ASVs were compared to trimmed or short-read sequences (Figure 8A–B). For the long- and trimmed-read comparison (Figure 8A), agreement was higher at broader levels, with only 29–32% of ASVs showing mismatched assignments from the domain to the subdivision. Discrepancies increased with taxonomic resolution, reaching approximately 39% at the family level and nearly 49% at the genus and species levels. Similarly, when short-read ASVs were compared with long-read ASVs using BLAST, sequences sharing at least 98% base pair identity were used to evaluate taxonomic consistency (Figure 8B). Here, mismatches were negligible at higher levels (<2%) and remained low up to the order level (≤9%), but increased sharply at finer ranks, reaching 12.7%, 37.4 %, and 47.9% at the family, genus, and species levels, respectively.

**Figure 8.**
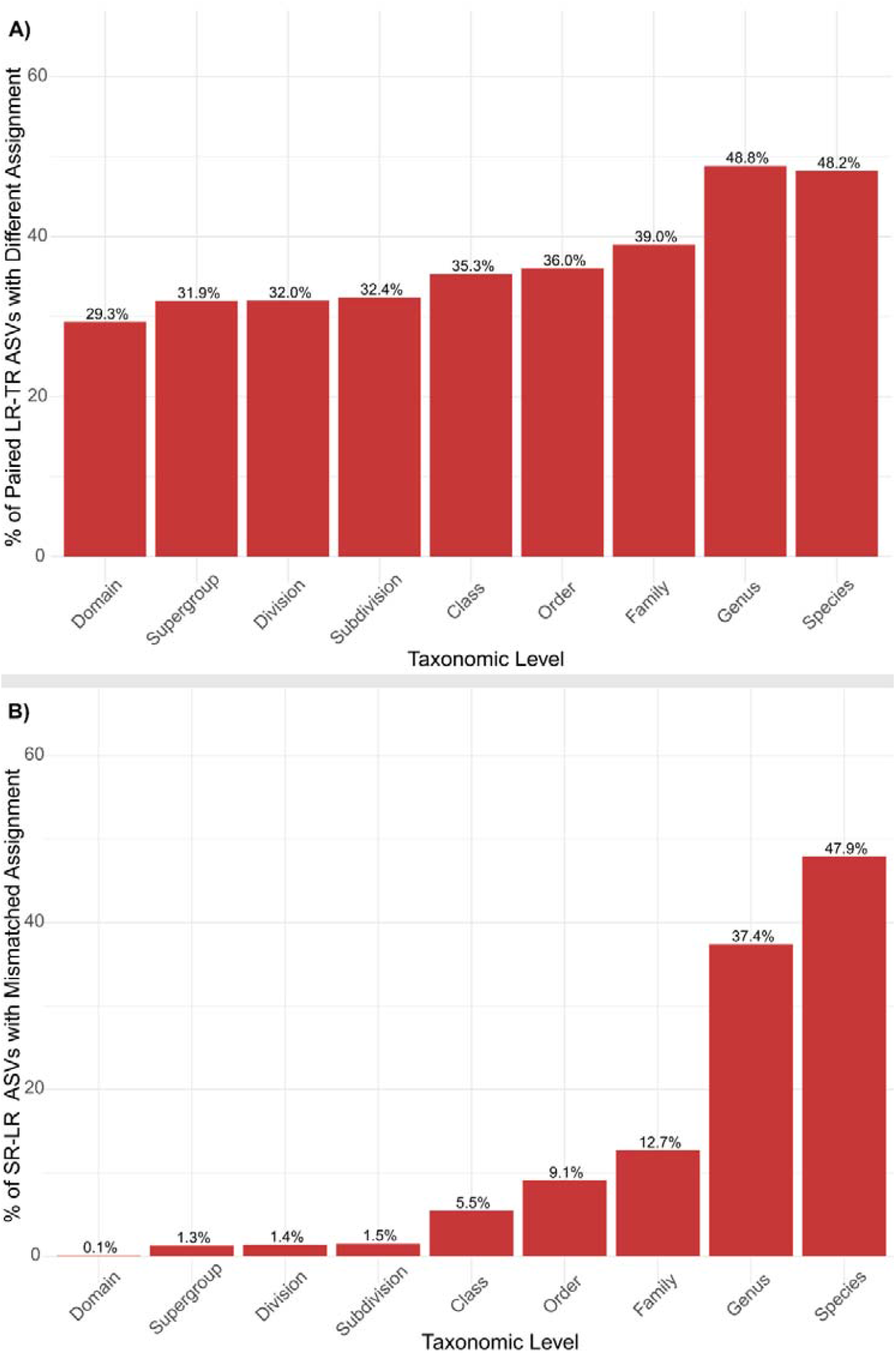
Comparison of taxonomic agreement between sequencing approaches. (A) Percentage of paired long-read (LR) and trimmed-read (TR) ASVs with different taxonomic assignments across ranks. (B) Comparison between short-read (SR) and long-read (LR) ASVs, where SR sequences were matched to LR ASVs using BLAST, and only those with ≥98% sequence identity were retained.

Together, these results indicate that while higher-level classifications are broadly consistent across sequencing types, lower-level taxonomic resolution, particularly at the genus and species levels, shows greater variability. Notably, the 137 species unique to the trimmed dataset and 161 species unique to the short-read dataset (Figure 7B) may, in part, reflect such discrepancies, as nearly half of the paired long–read and trimmed ASVs differed in their species assignment, similar to that of the paired short-read and long-read ASVs. This suggests that some of the apparent “unique” taxa in the trimmed and short-read datasets likely result from differences in classification accuracy rather than the detection of genuinely distinct species.

## Discussion

Rivers are highly dynamic ecosystems in which physical, chemical, and biological gradients shape complex microbial and algal communities (Battin et al., 2016). Within these environments, biofilms represent ecologically significant assemblages whose composition responds sensitively to environmental variation and pollution gradients (Kelly *et al*., 2020). Therefore, the accurate characterisation of these communities is critical for ecological monitoring and water quality assessment. Molecular approaches based on eDNA have become invaluable tools for identifying microbial assemblages. However, methodological factors, such as sequencing technology, read length and primer choice, can markedly influence taxonomic resolution and the interpretation of bacterial biofilm diversity patterns (Anderson *et al*., 2025). In this study, we compared long- and short-read sequencing data, as well as trimmed long-read data that mimicked the short-read region, to assess the impact of read length and sequencing platform on microeukaryote (18S) community profiling.

Our findings demonstrate clear differences between sequencing methods in terms of taxonomic assignment and diversity estimates. Unlike our previous analysis of bacterial 16S rRNA genes (Anderson *et al*., 2025), where sequencing platform had minimal influence on community composition, the choice of sequencing technology for 18S metabarcoding had a pronounced effect. The long-read dataset displayed a higher overall taxonomic resolution, consistent with previous reports highlighting the enhanced discriminatory power of long-read approaches for eukaryotic metabarcoding in other environments (Krehenwinkel *et al*., 2020; Bylemans *et al*., 2025). The increased assignment rates and finer-level classifications indicate that full-length amplicons capture more variable regions of the marker gene, thereby reducing the ambiguity in database matching. Importantly, however, our comparative analysis of the trimmed dataset revealed that shortening long reads to match the short-read target region significantly reduced taxonomic precision and led to greater inconsistencies at lower taxonomic ranks. Nearly half of the paired long-read and trimmed ASVs differed in genus or species assignment (Figure 8A), despite representing the same underlying sequence variants. This suggests that the apparent “unique” taxa in the trimmed dataset (and by extension, in short-read data) may be artefacts of misclassification rather than genuinely distinct species.

Further supporting this interpretation, our BLAST-based comparison of short-read ASVs to their long-read counterparts (Figure 8B) demonstrated that mismatches were negligible at broad taxonomic levels (<2%) but increased substantially at finer resolutions, reaching nearly 48% at the species level. These discrepancies are likely driven by the limited discriminatory power of shorter reads, which contain fewer diagnostic positions for distinguishing between closely related taxa. Consequently, although short-read sequencing provides reliable community overviews at higher ranks, it may misrepresent species-level diversity in complex assemblages. The trimmed dataset in particular serves as an informative proxy, illustrating how reduced read length alone can lead to systematic shifts in taxonomic assignments, even when sequencing accuracy is held constant.

The influence of read length on taxonomic concordance has important implications for biomonitoring applications. Although long-read platforms offer greater taxonomic depth and fewer unclassified ASVs, they remain more costly and currently yield lower throughput than short-read platforms such as Illumina (Tedersoo et al., 2021). Therefore, the optimal sequencing strategy depends on the ecological question and taxonomic resolution required. For large-scale monitoring programmes where genus- or family-level data are sufficient, short-read sequencing remains cost-effective and reproducible. However, for studies requiring accurate species level identification, such as assessments of invasive or protected species, long-read sequencing provides a critical advantage. As observed in this study, the apparent species richness derived from short-read data must be interpreted cautiously, as shorter amplicons can inflate diversity metrics by introducing misclassified variants.

In addition to read length, primer choice is a well-recognised source of bias in 18S metabarcoding studies and may also influence observed community composition. Previous work has demonstrated that primer degeneracy, GC content sensitivity, and target region selection can alter relative abundance estimates and reduce detection of certain taxa, particularly when using short V4 amplicons (Bradley et al., 2016; Wylezich et al., 2017). Comparative evaluations of full-length and short-read 18S markers have similarly reported lower genus-level recovery and reduced taxonomic resolution when targeting short regions alone (Latz et al., 2022; Gaonkar & Campbell, 2018). In the present study, the inclusion of a trimmed long-read dataset allowed partial isolation of read-length effects from platform-specific biases, as the same sequence variants were evaluated across differing amplicon lengths. While primer-region targeting likely contributes to differences between datasets, the high concordance between full-length and trimmed long reads at the community level, combined with increasing taxonomic mismatches at finer ranks, indicates that reduced sequence length itself remains a primary driver of classification inconsistency. These findings are consistent with recommendations that primer standardisation and careful marker selection are essential for cross-study comparability in eDNA-based monitoring (Lear et al., 2018).

Our results also highlight the need for continued refinement of freshwater 18S rRNA gene reference databases. Even with the increased resolution afforded by long-read sequencing, some ASVs could not be confidently assigned to species, reflecting ongoing gaps in the curated freshwater taxon records. Future efforts to expand reference coverage and improve taxonomic curation are essential to fully realise the potential of long-read metabarcoding for ecological monitoring. Improved database standardisation will also enhance cross-platform comparability, reducing inconsistencies, such as those observed here between sequencing methods.

Collectively, this study demonstrates that sequencing methods exert a measurable influence on the inferred freshwater community structure. While both long- and short-read approaches detected comparable dominant taxa, they did not capture broadly consistent beta-diversity patterns, with each method producing statistically distinct community profiles. The long-read and trimmed long-read datasets showed near-identical compositions, confirming that trimming long reads to match short-read primer regions does not distort the overall community patterns. In contrast, short-read sequencing generates markedly different assemblage structures, likely reflecting methodological biases in taxon recovery or abundance estimation. The trimmed dataset further revealed that apparent “unique” taxa may arise from misclassification linked to reduced sequence length, rather than true biological differences. These findings emphasise that read length and sequencing strategy strongly influence ecological interpretations derived from eDNA metabarcoding using the 18S rRNA gene. Consequently, careful consideration of sequencing technology and target regions is essential when designing biomonitoring studies, particularly where fine-scale taxonomic resolution is required.

## Authors contribution

Physicochemical data and samples were collected by the Environment Agency. D.S.R., M.A., S.B.B., K.W., and J.W. conceptualised the study. M.A. and A.C.T performed sample processing, DNA extractions and sequencing. M.A. and S.B.B. designed and performed the bioinformatic analyses. M.A. performed statistical analysis and figure production. M.A. drafted the manuscript. All authors have read and revised the manuscript.

## Data Accessibility statement

Raw sequence reads and related metadata can be found at:

