## Supplementary material for "Resolving eukaryotic river biofilm communities using long-read sequencing for biomonitoring"

Supplementary Table 1: Primers used in this study.

| Sequencing Method | Forward primer (5’-3’) | Reverse primer (5’-3’) | Reference |
| --- | --- | --- | --- |
| Illumina 1st round | NSF573:  CGCGGTAATTCCAGCTCCA | NSR951:  TTGGYRAATGCTTTCGC | Mangot et al., 2012 |
| Illumina 1st round adaptor | Forward adaptor: TCGTCGGCAGCGTCAGATGTGTATAAGAGAC | Reverse adaptor: GTCTCGTGGGCTCGGAGATGTGTATAAGAGACAG |  |
| Pacific Biosciences | Euk-A 18S V1-V9:  AACCTGGTTGATCCTGCCAGT | Euk-B 18S V1-V9:  GATCCTTCTGCAGGTTCACCTAC | Countway et al., 2005 |

Supplementary Table 2: PCR conditions.

| Sequencing Method | Stage | Temperature °C | Time | Number of cycles |
| --- | --- | --- | --- | --- |
| Illumina 1st round | Initial denaturation  Denaturation  Annealing  Extension  Final extension | 94  94  60  72  72 | 5 min  30 sec  30 sec  30 sec  10 min | 30 |
| Illumina  Indexing | Initial denaturation  Denaturation  Annealing  Extension  Final extension | 95  95  50  72  72 | 2 min  15 sec  30 sec  30 sec  10 min | 8 |
| Pacific Biosciences | Initial denaturation  Denaturation  Annealing  Extension  Final extension | 95  95  50  72  72 | 2 min  30 sec  30 sec  2.5 min  7 min | 35 |

Supplementary Table 4: P values from Kruskal Wallis test to test for significant differences in taxonomic assignment between sequencing groups at each taxonomic level.

| Taxonomic_Level | group1 | group2 | p | p.adj | p.adj.signif |
| --- | --- | --- | --- | --- | --- |
| Supergroup | Long | Short | 1.6e-05 | 2.4e-05 | **** |
| Supergroup | Long | Trimmed | 5.12e-09 | 1.54e-08 | **** |
| Supergroup | Short | Trimmed | 0.001 | 0.001 | ** |
| Division | Long | Short | 0.000364 | 0.000546 | *** |
| Division | Long | Trimmed | 1.79e-08 | 5.37e-08 | **** |
| Division | Short | Trimmed | 0.000741 | 0.000741 | *** |
| Subdivision | Long | Short | 0.000262 | 0.000262 | *** |
| Subdivision | Long | Trimmed | 9.87e-09 | 2.96e-08 | **** |
| Subdivision | Short | Trimmed | 4.17e-05 | 6.26e-05 | **** |
| Class | Long | Short | 0.004 | 0.004 | ** |
| Class | Long | Trimmed | 2.07e-08 | 6.21e-08 | **** |
| Class | Short | Trimmed | 7.91e-07 | 1.19e-06 | **** |
| Order | Long | Short | 0.105 | 0.105 | ns |
| Order | Long | Trimmed | 4.04e-08 | 1.21e-07 | **** |
| Order | Short | Trimmed | 7.66e-07 | 1.15e-06 | **** |
| Family | Long | Short | 0.004 | 0.004 | ** |
| Family | Long | Trimmed | 1.03e-09 | 3.09e-09 | **** |
| Family | Short | Trimmed | 7.3e-09 | 1.1e-08 | **** |
| Genus | Long | Short | 6.5e-09 | 9.75e-09 | **** |
| Genus | Long | Trimmed | 1.84e-10 | 5.52e-10 | **** |
| Genus | Short | Trimmed | 0.024 | 0.024 | * |
| Species | Long | Short | 0.001 | 0.001 | ** |
| Species | Long | Trimmed | 5.02e-08 | 1.51e-07 | **** |
| Species | Short | Trimmed | 7.39e-06 | 1.11e-05 | **** |

Supplementary Table 5: Relative abundance for each sequencing type at the Division level.

| Taxonomy | Sequencing  type | Abundance | Standard deviation | Standard  error |
| --- | --- | --- | --- | --- |
| Actinobacteria | Long | 0.014583 | 0.059189375 | 0.009133119 |
| Actinobacteria | Short | 0.064269 | 0.168429407 | 0.025989222 |
| Actinobacteria | Trimmed | 0.000000 | 0 | 0 |
| Alveolata | Long | 11.960670 | 14.12922469 | 2.180186703 |
| Alveolata | Short | 6.128319 | 8.288635518 | 1.278964227 |
| Alveolata | Trimmed | 10.089230 | 13.00913847 | 2.007353646 |
| Amoebozoa_X | Long | 0.001267 | 0.008214179 | 0.001267475 |
| Amoebozoa_X | Short | 0.000000 | 0 | 0 |
| Amoebozoa_X | Trimmed | 0.000000 | 0 | 0 |
| Ancyromonadida | Long | 0.002606 | 0.010114321 | 0.001560674 |
| Ancyromonadida | Short | 0.002826 | 0.007937537 | 0.001224789 |
| Ancyromonadida | Trimmed | 0.005079 | 0.018241916 | 0.002814789 |
| Apusomonada | Long | 0.019985 | 0.054724895 | 0.008444235 |
| Apusomonada | Short | 0.000169 | 0.001094919 | 0.00016895 |
| Apusomonada | Trimmed | 0.006293 | 0.023216887 | 0.003582443 |
| Breviatea | Long | 0.005079 | 0.029516134 | 0.004554438 |
| Breviatea | Short | 0.001077 | 0.005458797 | 0.000842311 |
| Breviatea | Trimmed | 0.006287 | 0.033458754 | 0.005162798 |
| Centroplasthelida | Long | 0.025524 | 0.055821574 | 0.008613456 |
| Centroplasthelida | Short | 0.000000 | 0 | 0 |
| Centroplasthelida | Trimmed | 0.031629 | 0.047226476 | 0.007287204 |
| Chlorophyta | Long | 8.408230 | 8.831794946 | 1.362775546 |
| Chlorophyta | Short | 12.144050 | 13.59204408 | 2.097297934 |
| Chlorophyta | Trimmed | 9.359665 | 9.682280412 | 1.494008303 |
| Cryptophyta | Long | 0.211996 | 0.695201774 | 0.107271963 |
| Cryptophyta | Short | 0.039932 | 0.161795008 | 0.024965512 |
| Cryptophyta | Trimmed | 0.228521 | 0.728375623 | 0.112390799 |
| Cryptophyta:nucl | Long | 0.002408 | 0.011160329 | 0.001722076 |
| Cryptophyta:nucl | Short | 0.000000 | 0 | 0 |
| Cryptophyta:nucl | Trimmed | 0.003335 | 0.015316239 | 0.002363347 |
| Discoba | Long | 0.013617 | 0.037880715 | 0.005845121 |
| Discoba | Short | 0.000000 | 0 | 0 |
| Discoba | Trimmed | 0.000000 | 0 | 0 |
| Discosea | Long | 0.046178 | 0.067412137 | 0.010401919 |
| Discosea | Short | 0.009814 | 0.019205567 | 0.002963483 |
| Discosea | Trimmed | 0.010898 | 0.019518194 | 0.003011723 |
| Evosea | Long | 0.088811 | 0.262831957 | 0.040555851 |
| Evosea | Short | 0.000713 | 0.004622167 | 0.000713216 |
| Evosea | Trimmed | 0.084881 | 0.246634611 | 0.038056547 |
| Haptophyta | Long | 0.000000 | 0 | 0 |
| Haptophyta | Short | 0.004230 | 0.017046141 | 0.002630277 |
| Haptophyta | Trimmed | 0.001215 | 0.005783404 | 0.000892399 |
| Kathablepharidacea | Long | 0.002203 | 0.008258878 | 0.001274373 |
| Kathablepharidacea | Short | 0.003628 | 0.011381172 | 0.001756153 |
| Kathablepharidacea | Trimmed | 0.004646 | 0.012798465 | 0.001974846 |
| Metamonada | Long | 0.000000 | 0 | 0 |
| Metamonada | Short | 0.000364 | 0.002358928 | 0.00036399 |
| Metamonada | Trimmed | 0.000000 | 0 | 0 |
| Nibbleridia | Long | 0.000000 | 0 | 0 |
| Nibbleridia | Short | 0.000268 | 0.001735826 | 0.000267844 |
| Nibbleridia | Trimmed | 0.000000 | 0 | 0 |
| Opisthokonta | Long | 34.272190 | 24.99173465 | 3.856308378 |
| Opisthokonta | Short | 44.283670 | 24.38605663 | 3.762850231 |
| Opisthokonta | Trimmed | 34.671210 | 23.87073331 | 3.683334115 |
| Prasinodermophyta | Long | 0.003531 | 0.016008764 | 0.002470206 |
| Prasinodermophyta | Short | 0.000000 | 0 | 0 |
| Prasinodermophyta | Trimmed | 0.000000 | 0 | 0 |
| Rhizaria | Long | 4.387355 | 6.354220853 | 0.980477564 |
| Rhizaria | Short | 3.287070 | 4.048953952 | 0.624767159 |
| Rhizaria | Trimmed | 4.827898 | 5.988921547 | 0.924110657 |
| Rhodophyta | Long | 12.208560 | 18.91058712 | 2.917966943 |
| Rhodophyta | Short | 0.001936 | 0.007075436 | 0.001091764 |
| Rhodophyta | Trimmed | 11.984610 | 18.6208565 | 2.873260537 |
| Rigifilida | Long | 0.008035 | 0.038546045 | 0.005947784 |
| Rigifilida | Short | 0.059203 | 0.224881902 | 0.034700031 |
| Rigifilida | Trimmed | 0.010609 | 0.034065397 | 0.005256405 |
| Stramenopiles | Long | 22.561520 | 23.6560987 | 3.650215276 |
| Stramenopiles | Short | 22.879870 | 19.91402299 | 3.072800459 |
| Stramenopiles | Trimmed | 22.232310 | 22.71557873 | 3.505089894 |
| Streptophyta | Long | 5.441885 | 13.12236822 | 2.024825376 |
| Streptophyta | Short | 10.548500 | 14.58131126 | 2.249945175 |
| Streptophyta | Trimmed | 5.370953 | 12.49264717 | 1.927657309 |
| Telonemia | Long | 0.000000 | 0 | 0 |
| Telonemia | Short | 0.000225 | 0.001459893 | 0.000225266 |
| Telonemia | Trimmed | 0.001624 | 0.00605932 | 0.000934973 |
| Tubulinea | Long | 0.109373 | 0.168971147 | 0.026072814 |
| Tubulinea | Short | 0.041981 | 0.091249485 | 0.014080101 |
| Tubulinea | Trimmed | 0.119734 | 0.144501808 | 0.022297113 |
| unidentified | Long | 0.204384 | 0.342469037 | 0.05284412 |
| unidentified | Short | 0.497879 | 0.832260943 | 0.128420652 |
| unidentified | Trimmed | 0.949364 | 1.112109458 | 0.171602215 |

Supplementary Table 6: Relative abundance for each sequencing type at the for the top 50 species.

| Taxonomy | Sample | Abundance | SD | SE | mean  abundance |
| --- | --- | --- | --- | --- | --- |
| unidentified | Long | 30.39752273 | 18.22736447 | 2.81E+00 | 0.412819466 |
| unidentified | Short | 45.30074416 | 19.77870039 | 3.05E+00 | 0.412819466 |
| unidentified | Trimmed | 48.14757298 | 18.51095157 | 2.86E+00 | 0.412819466 |
| Sirodotia_delicatula | Long | 7.764815383 | 10.97950528 | 1.69E+00 | 0.05129716 |
| Sirodotia_delicatula | Short | 0.00133873 | 0.006060321 | 9.35E-04 | 0.05129716 |
| Sirodotia_delicatula | Trimmed | 7.6229939 | 10.74516137 | 1.66E+00 | 0.05129716 |
| Navicula_tripunctata | Long | 2.719731322 | 6.018988963 | 9.29E-01 | 0.023831305 |
| Navicula_tripunctata | Short | 1.87348774 | 3.242765225 | 5.00E-01 | 0.023831305 |
| Navicula_tripunctata | Trimmed | 2.556172344 | 5.701035534 | 8.80E-01 | 0.023831305 |
| Brachypodium_distachyon | Long | 2.30751568 | 11.2428958 | 1.73E+00 | 0.020769532 |
| Brachypodium_distachyon | Short | 2.120483672 | 10.70427719 | 1.65E+00 | 0.020769532 |
| Brachypodium_distachyon | Trimmed | 1.802860379 | 10.07974089 | 1.56E+00 | 0.020769532 |
| Nais_elinguis | Long | 0.089187466 | 0.472598606 | 7.29E-02 | 0.018135388 |
| Nais_elinguis | Short | 3.459187628 | 8.812579863 | 1.36E+00 | 0.018135388 |
| Nais_elinguis | Trimmed | 1.892241343 | 6.400705186 | 9.88E-01 | 0.018135388 |
| Balbiania_investiens | Long | 2.404766872 | 6.396523439 | 9.87E-01 | 0.015920077 |
| Balbiania_investiens | Short | 0 | 0 | 0.00E+00 | 0.015920077 |
| Balbiania_investiens | Trimmed | 2.371256114 | 6.274432235 | 9.68E-01 | 0.015920077 |
| Hydrurus_foetidus | Long | 1.17441881 | 4.585809245 | 7.08E-01 | 0.015222193 |
| Hydrurus_foetidus | Short | 2.292635061 | 10.03065837 | 1.55E+00 | 0.015222193 |
| Hydrurus_foetidus | Trimmed | 1.099604131 | 4.290128031 | 6.62E-01 | 0.015222193 |
| Mattesia_geminata | Long | 2.20448638 | 11.85466511 | 1.83E+00 | 0.014355923 |
| Mattesia_geminata | Short | 0 | 0 | 0.00E+00 | 0.014355923 |
| Mattesia_geminata | Trimmed | 2.102290564 | 11.56010604 | 1.78E+00 | 0.014355923 |
| Melosira_varians | Long | 1.955770097 | 6.074593262 | 9.37E-01 | 0.013318945 |
| Melosira_varians | Short | 0.177667804 | 0.623808087 | 9.63E-02 | 0.013318945 |
| Melosira_varians | Trimmed | 1.862245687 | 5.723267664 | 8.83E-01 | 0.013318945 |
| Polycelis_tenuis | Long | 3.79188296 | 13.23324219 | 2.04E+00 | 0.01263961 |
| Polycelis_tenuis | Short | 0 | 0 | 0.00E+00 | 0.01263961 |
| Polycelis_tenuis | Trimmed | 0 | 0 | 0.00E+00 | 0.01263961 |
| Cryptococcus_carnescens | Long | 1.767127795 | 6.199502786 | 9.57E-01 | 0.01230357 |
| Cryptococcus_carnescens | Short | 0.251122464 | 0.673405566 | 1.04E-01 | 0.01230357 |
| Cryptococcus_carnescens | Trimmed | 1.672820635 | 5.509289863 | 8.50E-01 | 0.01230357 |
| Cladophora_glomerata | Long | 0 | 0 | 0.00E+00 | 0.012112742 |
| Cladophora_glomerata | Short | 3.181236991 | 8.511258535 | 1.31E+00 | 0.012112742 |
| Cladophora_glomerata | Trimmed | 0.452585584 | 1.25730139 | 1.94E-01 | 0.012112742 |
| Chlorochytrium_lemnae | Long | 1.439485416 | 4.846282483 | 7.48E-01 | 0.011824944 |
| Chlorochytrium_lemnae | Short | 0.836605373 | 2.756811388 | 4.25E-01 | 0.011824944 |
| Chlorochytrium_lemnae | Trimmed | 1.271392482 | 3.828750687 | 5.91E-01 | 0.011824944 |
| Ulothrix_zonata | Long | 0.044099905 | 0.264069303 | 4.07E-02 | 0.011261342 |
| Ulothrix_zonata | Short | 2.287542966 | 7.701656095 | 1.19E+00 | 0.011261342 |
| Ulothrix_zonata | Trimmed | 1.046759802 | 3.415414863 | 5.27E-01 | 0.011261342 |
| Diatoma_tenue | Long | 1.540742463 | 7.637524394 | 1.18E+00 | 0.011183809 |
| Diatoma_tenue | Short | 0.287496146 | 1.021159348 | 1.58E-01 | 0.011183809 |
| Diatoma_tenue | Trimmed | 1.526904058 | 7.57847716 | 1.17E+00 | 0.011183809 |
| Nais_communis | Long | 2.63243458 | 7.986427233 | 1.23E+00 | 0.010159976 |
| Nais_communis | Short | 0.350831482 | 1.347579147 | 2.08E-01 | 0.010159976 |
| Nais_communis | Trimmed | 0.064726735 | 0.220034441 | 3.40E-02 | 0.010159976 |
| Lecythium_hyalinum | Long | 0.952177917 | 2.615176589 | 4.04E-01 | 0.010096325 |
| Lecythium_hyalinum | Short | 1.115938478 | 2.925525756 | 4.51E-01 | 0.010096325 |
| Lecythium_hyalinum | Trimmed | 0.960781071 | 2.512329944 | 3.88E-01 | 0.010096325 |
| Moesziomyces_antarcticus | Long | 2.904319725 | 13.1200487 | 2.02E+00 | 0.009681066 |
| Moesziomyces_antarcticus | Short | 0 | 0 | 0.00E+00 | 0.009681066 |
| Moesziomyces_antarcticus | Trimmed | 0 | 0 | 0.00E+00 | 0.009681066 |
| Sanionia_uncinata | Long | 0 | 0 | 0.00E+00 | 0.009639097 |
| Sanionia_uncinata | Short | 2.124478752 | 5.890379703 | 9.09E-01 | 0.009639097 |
| Sanionia_uncinata | Trimmed | 0.767250261 | 2.019432026 | 3.12E-01 | 0.009639097 |
| Vorticella_campanula | Long | 0.464612921 | 1.359439901 | 2.10E-01 | 0.008385534 |
| Vorticella_campanula | Short | 1.600039587 | 5.379842476 | 8.30E-01 | 0.008385534 |
| Vorticella_campanula | Trimmed | 0.451007682 | 1.274959428 | 1.97E-01 | 0.008385534 |
| Uncinais_uncinata | Long | 0 | 0 | 0.00E+00 | 0.008041818 |
| Uncinais_uncinata | Short | 1.869248694 | 4.167985988 | 6.43E-01 | 0.008041818 |
| Uncinais_uncinata | Trimmed | 0.543296747 | 1.36907021 | 2.11E-01 | 0.008041818 |
| Salmo_salar | Long | 0.301906638 | 1.939930302 | 2.99E-01 | 0.007250721 |
| Salmo_salar | Short | 1.172692198 | 7.553109331 | 1.17E+00 | 0.007250721 |
| Salmo_salar | Trimmed | 0.70061751 | 4.404627033 | 6.80E-01 | 0.007250721 |
| Orthocladius_luteipes | Long | 0 | 0 | 0.00E+00 | 0.007186294 |
| Orthocladius_luteipes | Short | 2.155888211 | 7.428606969 | 1.15E+00 | 0.007186294 |
| Orthocladius_luteipes | Trimmed | 0 | 0 | 0.00E+00 | 0.007186294 |
| Navicula_gregaria | Long | 0.723393025 | 1.792227921 | 2.77E-01 | 0.007177188 |
| Navicula_gregaria | Short | 0.769861001 | 1.72565635 | 2.66E-01 | 0.007177188 |
| Navicula_gregaria | Trimmed | 0.659902516 | 1.617370929 | 2.50E-01 | 0.007177188 |
| Cocconeis_placentula | Long | 0.05364954 | 0.228543852 | 3.53E-02 | 0.007066844 |
| Cocconeis_placentula | Short | 1.993402747 | 3.969825768 | 6.13E-01 | 0.007066844 |
| Cocconeis_placentula | Trimmed | 0.073000977 | 0.245585199 | 3.79E-02 | 0.007066844 |
| Polycelis_felina | Long | 0.610284265 | 2.132786798 | 3.29E-01 | 0.006876772 |
| Polycelis_felina | Short | 0.884927213 | 2.844600811 | 4.39E-01 | 0.006876772 |
| Polycelis_felina | Trimmed | 0.567820087 | 1.990447723 | 3.07E-01 | 0.006876772 |
| Desmodesmus_pannonicus | Long | 1.457887834 | 2.724650662 | 4.20E-01 | 0.006352472 |
| Desmodesmus_pannonicus | Short | 0.116095667 | 0.661584378 | 1.02E-01 | 0.006352472 |
| Desmodesmus_pannonicus | Trimmed | 0.331758009 | 0.476614474 | 7.35E-02 | 0.006352472 |
| Navicula_cryptotenella | Long | 0.970622288 | 2.400385316 | 3.70E-01 | 0.006219071 |
| Navicula_cryptotenella | Short | 0.390712962 | 0.69483175 | 1.07E-01 | 0.006219071 |
| Navicula_cryptotenella | Trimmed | 0.504386043 | 1.467681175 | 2.26E-01 | 0.006219071 |
| Planophila_laetevirens | Long | 0.984877138 | 2.493858047 | 3.85E-01 | 0.006212804 |
| Planophila_laetevirens | Short | 0.366500748 | 0.828671258 | 1.28E-01 | 0.006212804 |
| Planophila_laetevirens | Trimmed | 0.512463378 | 1.392420121 | 2.15E-01 | 0.006212804 |
| Amphora_pediculus | Long | 0.722572109 | 2.240480583 | 3.46E-01 | 0.006122678 |
| Amphora_pediculus | Short | 0.29709094 | 0.579401423 | 8.94E-02 | 0.006122678 |
| Amphora_pediculus | Trimmed | 0.817140298 | 2.118745183 | 3.27E-01 | 0.006122678 |
| Audouinella_hermannii | Long | 1.742945549 | 3.696129087 | 5.70E-01 | 0.005809818 |
| Audouinella_hermannii | Short | 0 | 0 | 0.00E+00 | 0.005809818 |
| Audouinella_hermannii | Trimmed | 0 | 0 | 0.00E+00 | 0.005809818 |
| Gomphonema_parvulum | Long | 0 | 0 | 0.00E+00 | 0.005724468 |
| Gomphonema_parvulum | Short | 1.17472141 | 3.916419754 | 6.04E-01 | 0.005724468 |
| Gomphonema_parvulum | Trimmed | 0.542619018 | 2.253009225 | 3.48E-01 | 0.005724468 |
| Chaetogaster_diaphanus | Long | 1.635581993 | 4.374516855 | 6.75E-01 | 0.00545194 |
| Chaetogaster_diaphanus | Short | 0 | 0 | 0.00E+00 | 0.00545194 |
| Chaetogaster_diaphanus | Trimmed | 0 | 0 | 0.00E+00 | 0.00545194 |
| Achnanthidium_minutissimum | Long | 0.667872506 | 1.323085814 | 2.04E-01 | 0.005443469 |
| Achnanthidium_minutissimum | Short | 0.321987289 | 0.679440922 | 1.05E-01 | 0.005443469 |
| Achnanthidium_minutissimum | Trimmed | 0.643180813 | 1.246958691 | 1.92E-01 | 0.005443469 |
| Ceratodon_purpureus | Long | 1.407951422 | 5.298467204 | 8.18E-01 | 0.005238119 |
| Ceratodon_purpureus | Short | 0.100655018 | 0.429244891 | 6.62E-02 | 0.005238119 |
| Ceratodon_purpureus | Trimmed | 0.062829332 | 0.285994062 | 4.41E-02 | 0.005238119 |
| Vaucheria_bursata | Long | 0.005791495 | 0.030037193 | 4.63E-03 | 0.004779581 |
| Vaucheria_bursata | Short | 1.419284114 | 7.112581144 | 1.10E+00 | 0.004779581 |
| Vaucheria_bursata | Trimmed | 0.008798714 | 0.047502713 | 7.33E-03 | 0.004779581 |
| Ancylus_fluviatilis | Long | 1.148865911 | 3.268043454 | 5.04E-01 | 0.003829553 |
| Ancylus_fluviatilis | Short | 0 | 0 | 0.00E+00 | 0.003829553 |
| Ancylus_fluviatilis | Trimmed | 0 | 0 | 0.00E+00 | 0.003829553 |
| Simulium_sanctipauli | Long | 0 | 0 | 0.00E+00 | 0.003692605 |
| Simulium_sanctipauli | Short | 1.107781527 | 4.315817361 | 6.66E-01 | 0.003692605 |
| Simulium_sanctipauli | Trimmed | 0 | 0 | 0.00E+00 | 0.003692605 |
| Hypsibius_dujardini | Long | 0.170017675 | 0.719140182 | 1.11E-01 | 0.003402304 |
| Hypsibius_dujardini | Short | 0.673310518 | 2.570309143 | 3.97E-01 | 0.003402304 |
| Hypsibius_dujardini | Trimmed | 0.177363149 | 0.771700706 | 1.19E-01 | 0.003402304 |
| Scenedesmus_armatus | Long | 0 | 0 | 0.00E+00 | 0.003062435 |
| Scenedesmus_armatus | Short | 0.261822167 | 1.637048398 | 2.53E-01 | 0.003062435 |
| Scenedesmus_armatus | Trimmed | 0.656908376 | 1.358176072 | 2.10E-01 | 0.003062435 |
| Monostroma_grevillei | Long | 0.220147452 | 0.418648002 | 6.46E-02 | 0.002708155 |
| Monostroma_grevillei | Short | 0.350756004 | 0.709647567 | 1.10E-01 | 0.002708155 |
| Monostroma_grevillei | Trimmed | 0.241543105 | 0.464960251 | 7.17E-02 | 0.002708155 |
| Encyonema_silesacum | Long | 0.271153802 | 0.733778163 | 1.13E-01 | 0.002528871 |
| Encyonema_silesacum | Short | 0.296013146 | 0.719262625 | 1.11E-01 | 0.002528871 |
| Encyonema_silesacum | Trimmed | 0.191494473 | 0.459604757 | 7.09E-02 | 0.002528871 |
| Sperchon_violaceus | Long | 0.209468902 | 0.493858089 | 7.62E-02 | 0.002520822 |
| Sperchon_violaceus | Short | 0.331278047 | 0.761856217 | 1.18E-01 | 0.002520822 |
| Sperchon_violaceus | Trimmed | 0.215499565 | 0.486281218 | 7.50E-02 | 0.002520822 |
| Verrucaria_dolosa | Long | 0 | 0 | 0.00E+00 | 0.002411031 |
| Verrucaria_dolosa | Short | 0.723309319 | 2.811986583 | 4.34E-01 | 0.002411031 |
| Verrucaria_dolosa | Trimmed | 0 | 0 | 0.00E+00 | 0.002411031 |
| Anteholosticha_monilata | Long | 0.474099585 | 1.330775234 | 2.05E-01 | 0.002336231 |
| Anteholosticha_monilata | Short | 0 | 0 | 0.00E+00 | 0.002336231 |
| Anteholosticha_monilata | Trimmed | 0.226769741 | 1.063160219 | 1.64E-01 | 0.002336231 |
| Ranunculus_sceleratus | Long | 0.698408503 | 3.869131506 | 5.97E-01 | 0.002328028 |
| Ranunculus_sceleratus | Short | 0 | 0 | 0.00E+00 | 0.002328028 |
| Ranunculus_sceleratus | Trimmed | 0 | 0 | 0.00E+00 | 0.002328028 |
| Punctodora_ratzeburgensis | Long | 0.281451415 | 1.117687727 | 1.72E-01 | 0.002320506 |
| Punctodora_ratzeburgensis | Short | 0.163944277 | 0.714125934 | 1.10E-01 | 0.002320506 |
| Punctodora_ratzeburgensis | Trimmed | 0.250756022 | 0.985726419 | 1.52E-01 | 0.002320506 |
| Chlorococcum_oleofaciens | Long | 0.027487283 | 0.102110343 | 1.58E-02 | 0.002279979 |
| Chlorococcum_oleofaciens | Short | 0.277782184 | 0.555519313 | 8.57E-02 | 0.002279979 |
| Chlorococcum_oleofaciens | Trimmed | 0.378724189 | 0.704998994 | 1.09E-01 | 0.002279979 |
| Cyclotella_meneghiniana | Long | 0.340020444 | 1.039518417 | 1.60E-01 | 0.002270081 |
| Cyclotella_meneghiniana | Short | 0 | 0 | 0.00E+00 | 0.002270081 |
| Cyclotella_meneghiniana | Trimmed | 0.341003806 | 1.015723313 | 1.57E-01 | 0.002270081 |
| Placopyrenium_canellum | Long | 0 | 0 | 0.00E+00 | 0.002255525 |
| Placopyrenium_canellum | Short | 0.676657582 | 2.215862246 | 3.42E-01 | 0.002255525 |
| Placopyrenium_canellum | Trimmed | 0 | 0 | 0.00E+00 | 0.002255525 |

Supplementary Table 7: Co-occurrence network structure across long-read, short-read, and trimmed datasets at the ASV level.

| **Sequencing type** | **Long** | **Short** | **Trim** |
| --- | --- | --- | --- |
| **Nodes** | 366 | 947 | 473 |
| **Edges** | 1071 | 8557 | 2687 |
| **Density** | 0.0160341 | 0.0191034 | 0.024071 |
| **Transitivity** | 0.5313071 | 0.4375808 | 0.416435 |
| **Average Transitivity** | 0.5402037 | 0.3847041 | 0.4139718 |
| **Modularity** | 0.8663067 | 0.7194276 | 0.7962736 |
| **Diameter** | 18 | 7 | 8 |
| **Assortativity** | 0.4454396 | 0.5775699 | 0.2763265 |
| **Mean Distance** | 0.2936196 | 0.1609791 | 0.1346859 |
| **Max Cliques** | 359 | 2754 | 995 |
| **Clusters** | 36 | 6 | 3 |

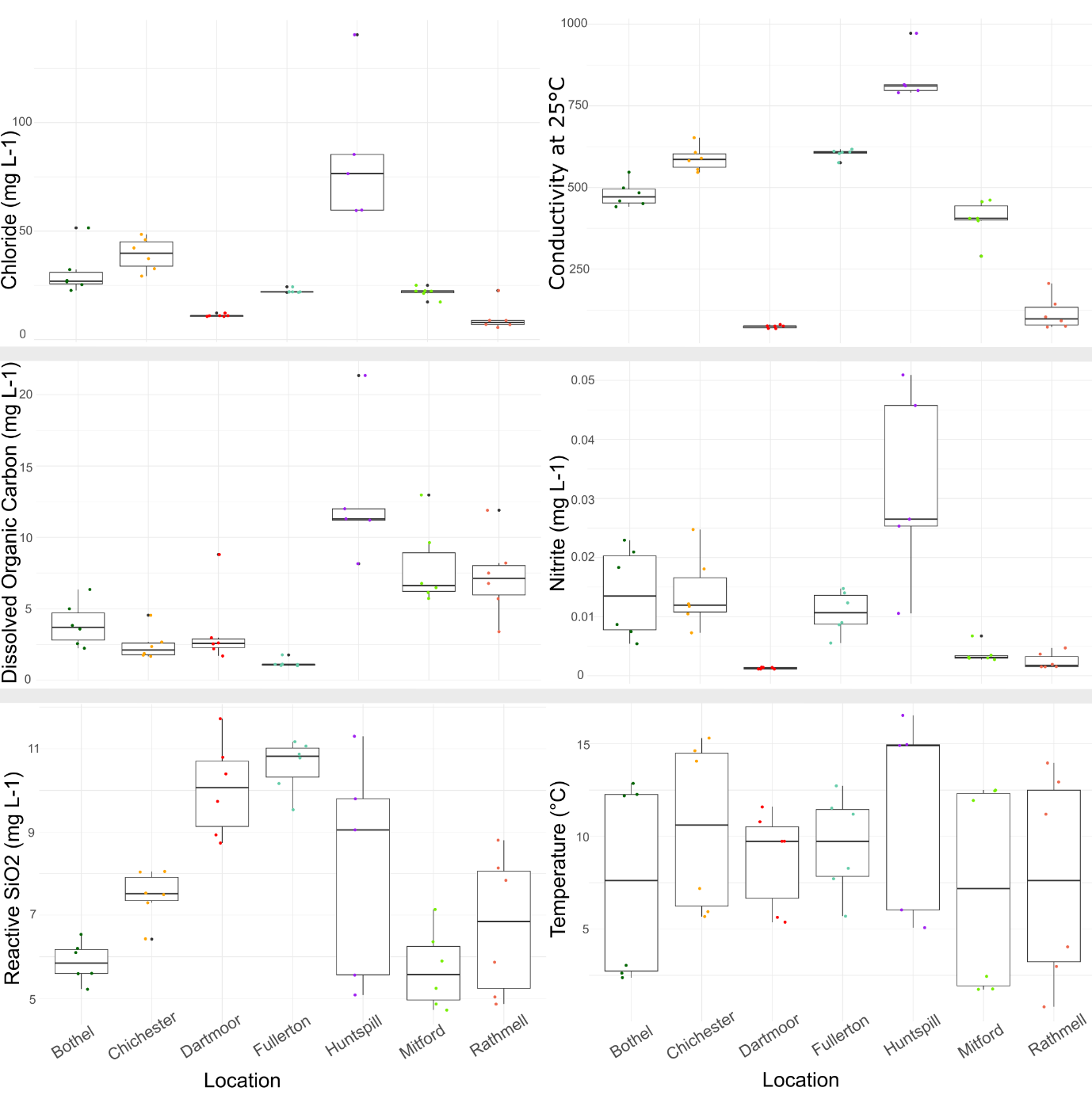

Supp Figure 1: chemistry data for the seven sampling sites; Chloride, Conductivity, Dissolved Organic Carbon, Nitrite, Reactive SiO2, and Temperature.

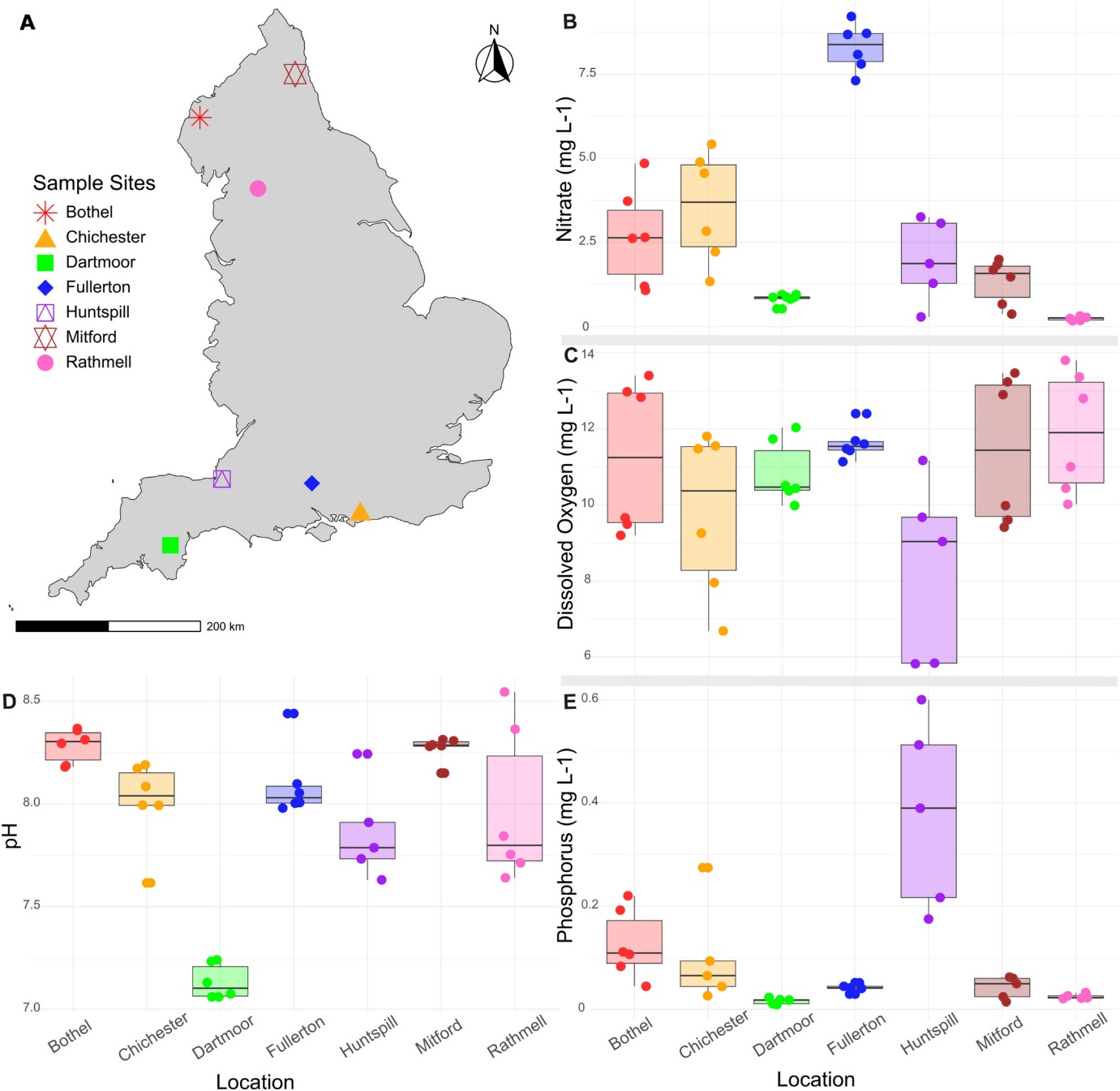

Supp Figure 2: (A) Map of the seven sites across England from which the biofilm samples were collected. Water chemistry data from the seven sites. Nitrate (mg L^−1^) (B), dissolved oxygen (mg L^−1^) (C), pH (D) and phosphorus (mg L^−1^) (E) levels at each sample point for each location. Water chemistry means were calculated using five chemistry samples collected over a three-month period prior to biofilm sample collection.

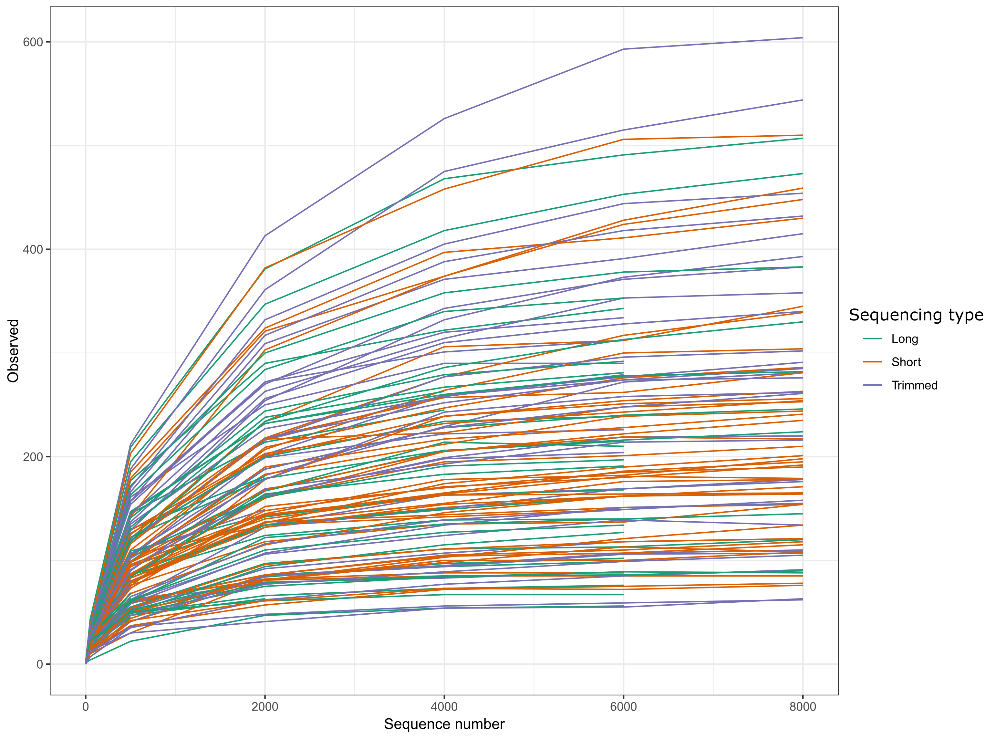

Supp Figure 3: Rarefaction curve for long-read, short-read and trimmed-read samples.

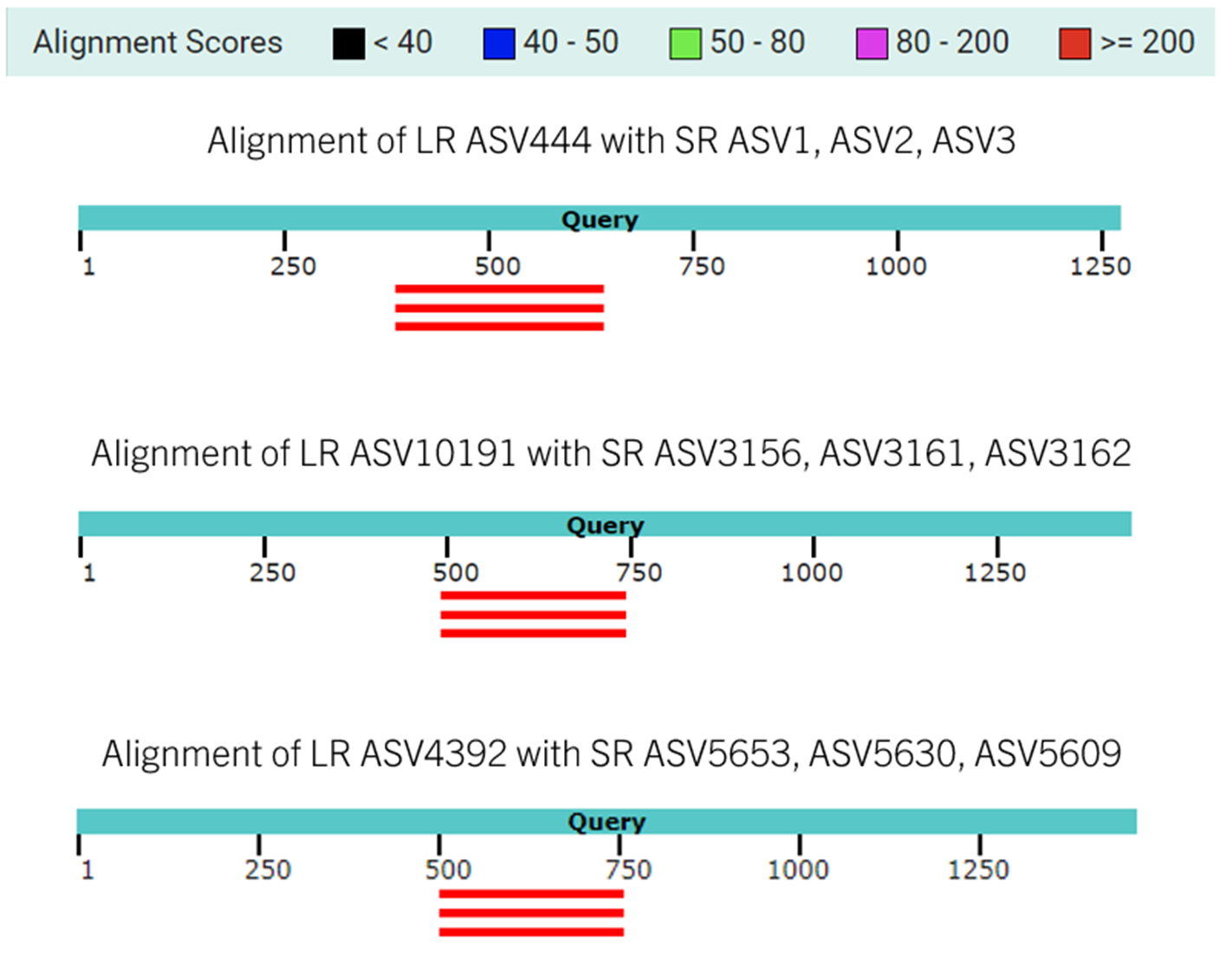

Supp Figure 4: Visualisation of BLAST alignments between long-read (LR) and short-read (SR) ASVs. The blue bar represents the LR ASV query sequence, while the red bars indicate the aligned SR ASVs that match the LR sequence. Alignment scores are colour-coded along the top bar, with red indicating a stronger alignment.
